# Mechanism of chaperone coordination during cotranslational protein folding in bacteria

**DOI:** 10.1101/2024.01.22.576655

**Authors:** Alžběta Roeselová, Sarah L. Maslen, Santosh Shivakumaraswamy, Grant A. Pellowe, Steven Howell, Dhira Joshi, Joanna Redmond, Svend Kjaer, J. Mark Skehel, David Balchin

## Abstract

Protein folding is assisted by molecular chaperones that bind nascent polypeptides during mRNA translation. Several structurally-distinct classes of chaperone promote *de novo* folding, suggesting that their activities are coordinated at the ribosome. We used biochemical reconstitution and structural proteomics to explore the molecular basis for cotranslational chaperone action in bacteria. We found that chaperone binding is disfavoured close to the ribosome, allowing folding to precede chaperone recruitment. Trigger factor subsequently recognises compact folding intermediates exposing extensive non-native surface and dictates DnaJ access to nascent chains. DnaJ uses a large surface to bind structurally diverse intermediates, and recruits DnaK to sequence-diverse solvent-accessible sites. Neither Trigger factor, DnaJ nor DnaK destabilize cotranslational folding intermediates. Instead, the chaperones collaborate to create a protected space for protein maturation that extends well beyond the ribosome exit tunnel. Our findings show how the chaperone network selects and modulates cotranslational folding intermediates.

## INTRODUCTION

Most newly synthesized polypeptides must fold into a specific three-dimensional conformation to function, and the failure to efficiently do so is a common cause of human disease ^1^. The complexity of protein folding is exacerbated for multidomain and oligomeric proteins, for which non-specific interdomain interactions and premature exposure of hydrophobic interfaces can hinder folding ^2,3^. *In vivo*, folding challenges are mitigated by the combined effect of cotranslational folding ^4,5^ and a network of molecular chaperones ^6^. These features of *de novo* folding are coupled, with recent work highlighting that chaperone recruitment to translating ribosomes is a general feature of protein biogenesis ^7,8^.

The chaperone network is largely conserved from bacteria to humans, and *E. coli* chaperones are paradigmatic examples. Although many chaperones bind nascent chains (NCs), the dominant interactors are Trigger factor (TF) and DnaJ/K (Hsp40/70 in eukaryotes) ^9^. TF and DnaJ/K share a subset of substrates and are at least partially redundant ^9–15^. Combined deletion of both chaperones leads to widespread aggregation of nascent proteins *in vivo* and is lethal under stress conditions ^15,16^. Extensive prior work has shown that TF can inhibit aggregation and delay NC folding, and in some cases may promote folding off the ribosome by outcompeting non-native long-range interactions ^17–24^. DnaK has primarily been studied in a post-translational context, where it has been demonstrated to bind hydrophobic motifs exposed prior to folding ^25^, protect partially-folded intermediates ^26–28^, and unfold misfolded substrates ^29–33^.

The folding function of chaperones is most commonly modelled using full-length client proteins that are artificially denatured or constitutively unfolded. However, during *de novo* folding, chaperones act on incompletely synthesized nascent polypeptides as they emerge from the ribosome exit tunnel. This scenario differs mechanistically from post-translational chaperone action in several respects. First, the folding intermediates encountered by chaperones during translation lack complete sequence information. Second, some chaperones have direct affinity for ribosome, which influences their access to nascent chains ^13,34^. Third, chaperone function at the ribosome occurs in the context of cotranslational folding. Vectorial synthesis, interactions with the ribosome surface, and sequestration of C-terminal sequences in the ribosome exit tunnel can all influence the stability and character of intermediates populated during *de novo* folding, especially for multidomain proteins ^4,5,19,35–47^. How molecular chaperones engage and process cotranslational folding intermediates is poorly understood.

Considering the ability of chaperones to interact promiscuously with non-native clients, a central question is how access to NCs is regulated. What features of NCs are recognised by different chaperones, and how is the binding of different chaperones coordinated? Due to the size and dynamic nature of chaperone complexes with partially folded nascent polypeptides, the mechanisms underpinning NC recognition by chaperones have remained elusive.

Here, we studied the interplay of molecular chaperones during cotranslational folding of the model multidomain protein β-galactosidase (β-gal). Using biochemical reconstitution, crosslinking-mass spectrometry and hydrogen/deuterium exchange-mass spectrometry, we characterised the interactions of ribosome-bound nascent β-gal with TF, DnaJ or DnaK, at different points throughout synthesis. Our results define the molecular basis for specific chaperone recruitment to nascent polypeptides and establish the structural consequences of chaperone binding.

## RESULTS

### Design of **β**-galactosidase ribosome:nascent chain complexes

As a model nascent chain we chose *Escherichia coli* β-gal, an endogenous chaperone substrate with a complex multidomain architecture (Fig 1A) ^17,48^. Each monomer of homotetrameric β-gal consists of 1023 aa and folds into 5 structured domains: a jelly roll β-barrel domain (D1), a TIM barrel-like domain (D3) containing the active site, two fibronectin type III barrel domains (D2 and D4) and a C-terminal β-sandwich (D5) ^49^. The N-terminal region is poorly structured. To sample different stages of β-gal synthesis we generated a series of stalled ribosome:nascent chain complexes (RNCs) in *E. coli* (Fig 1B). Translation was stalled at the point where each of the 5 individual domains had just emerged from the ribosome exit tunnel (assumed to protect ∼30 residues^50^), as well as multiple positions within each domain anticipated to preclude complete domain folding. RNC_1-1014_ mimics the state that would occur immediately prior to translation termination. As a negative control, we prepared RNC_40G/S_ in which the NC consists of a 40-residue Glycine/Serine-rich sequence. As a control for folded β-gal attached to the ribosome, we designed an RNC exposing the complete sequence of β-gal outside the exit tunnel via a C-terminal 50 aa linker (RNC_FL+50G/S_). Rigorous purification of RNCs under high-salt conditions yielded stable complexes containing intact ribosomes, with peptidyl tRNA-linked NCs at an abundance consistent with ∼100% occupancy (Fig 1C,D and S1A-D and Tables S1-3). RNC_FL+50G/S_ was enzymatically active and co-purified with excess β-gal chains without tRNA, consistent with oligomeric assembly into the native tetramer (Fig 1D and S1B,E). We used this set of RNCs, displaying structurally diverse cotranslational folding intermediates, as a platform to study chaperone interactions with nascent polypeptides at the ribosome.

**Figure 1.**
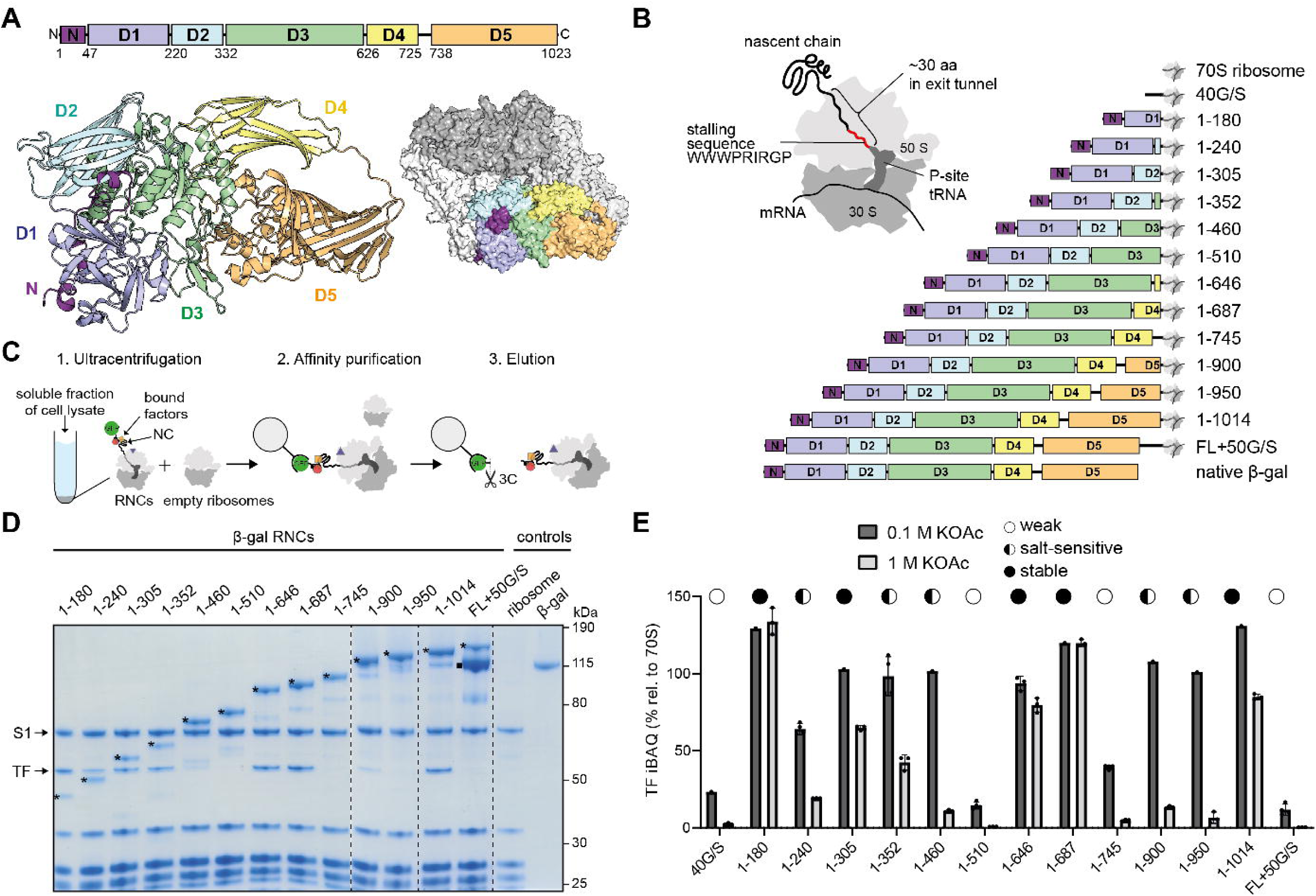
Stalled ribosome:nascent chain complexes selectively recruit Trigger factor. **(A)** Domain organisation and structure of *E. coli* β-galactosidase (β-gal). The monomer consists of an N-terminal extension (N, purple), jelly-roll β-barrel domain (D1, blue), a TIM barrel-like domain (D3, green), two fibronectin type III-like barrel domains (D2, cyan and D4, yellow) and a C-terminal β-sandwich (D5, orange). The structures of the β-gal monomer (left) and homotetramer (right) are coloured by domain (PDB: 6CVM^89^). **(B)** Schematic of truncated β-gal constructs encoded upstream of an arrest-enhanced variant of the SecM ribosome stalling sequence ^90^ (red). Domains are colour-coded as in (A). Artificial G/S-rich linker in RNC_40G/S_ and RNC_FL+50G/S_ is shown in black. **(C)** Ribosome:nascent chain complexe (RNC) purification scheme. The soluble fraction of cell lysates from *E. coli* overexpressing stalling constructs is centrifuged through a low-salt (0.1 M KOAc) or high-salt (1 M KOAc) sucrose cushion to isolate ribosomes (1). RNCs are separated from empty ribosomes via an affinity tag (muGFP) at the N-terminus of the NC (2). RNCs with any bound interactors are eluted by tag cleavage using HRV 3C (3). Optionally, RNCs are centrifuged through a second sucrose cushion. **(D)** Coomassie-stained SDS-PAGE of β-gal RNCs illustrated in (B), and purified as in (C) via two 1 M KOAc sucrose cushions. Bands corresponding to nascent chains (*) migrate higher (by ∼20 kDa) than expected based on protein molecular weight, due to the covalently bound tRNA. In FL+50G/S, released β-gal co-purifying with the RNC is indicated (▪). Some RNCs co-purify with Trigger factor (TF, ∼50 kDa). All other major bands correspond to 70S ribosomal proteins also present in the empty ribosome control. Except for S1 (∼70 kDa), all ribosomal proteins migrate below 40 kDa. The last lane contains purified full-length β-gal. **(E)** Intensity-based absolute quantification (iBAQ) of TF co-purified with RNCs under low (0.1 M KOAc) or high (1 M KOAc) salt conditions. Values are normalised to the average iBAQ value of all 70S ribosomal proteins. Depending on the RNC, TF co-purified either at low (○) or high (●) levels in both purification conditions, or exhibited salt-sensitive co-purification behaviour (IZl). Error-bars correspond to SD of 1-4 technical replicates. See also Fig S1 and Table S1-3.

### Selective recruitment of Trigger factor to nascent **β**-gal

We initially sought to identify endogenous NC-specific interactors that survive rigorous RNC purification. Mass spectrometry revealed the ribosome-associated molecular chaperone Trigger factor (TF) as the dominant co-purifying protein, abundant in several β-gal RNCs but absent in RNC_40G/S_ and RNC_FL+50G/S_ (Fig 1D and S1C, Table S1 and S2). As observed previously, we found that stable binding of TF to β-gal RNCs *in vitro* requires both ribosome and NC contacts, explaining the maximal occupancy of ∼1 TF per RNC (Fig S1F-I). The *K*_D_ for TF binding to RNC_1-646_ was ∼20 nM, as measured by quenching of BADAN-labelled TF fluorescence (Fig S1J) ^51^. This is similar to previous measurements with different TF substrates, and ∼100-fold lower than the *K*_D_ of ∼1-2 µM previously estimated for TF binding to empty ribosomes ^51–55^.

To understand how TF binding changes during β-gal synthesis we quantified co-purifying TF under low- and high-salt conditions (Fig 1E and S1K, Tables S1-3). At several NC lengths, TF occupancy remained above 50% even at 1 M KOAc, consistent with binding that is primarily stabilised by interactions with hydrophobic side chains expected to be exposed in incompletely folded NCs. In contrast, high salt stripped away the majority of TF from other RNCs, consistent with an alternative electrostatic binding mode. The folded control (RNC_FL+50G/S_) as well as two intermediate chain lengths – RNC_1-510_ (2½ domains) and RNC_1-745_ (4 domains) – were poorly recognized by TF in either salt condition. Notably, the mode of TF binding did not strictly correlate with domain emergence from the ribosome. Some NCs exposing complete domains (e.g. RNC_1-646_) engage TF via hydrophobic contacts, whereas incomplete domains in some NCs bury their hydrophobic surface from the chaperone (e.g. RNC_1-460_ and RNC_1-950_) or escape TF entirely (e.g. RNC_1-510_). The stability and character of the TF:NC interaction can therefore vary substantially during synthesis of a multidomain protein.

### The CTD and PPD of TF form a versatile NC-binding cavity

To probe the molecular basis of TF binding to NCs, we first analysed TF:RNC assemblies using crosslinking coupled to mass spectrometry (XL-MS), which provides residue-level distance restraints that report on the structural organisation of protein complexes. RNCs that co-purified with endogenous TF were crosslinked using the homobifunctional crosslinker disuccinimidyl dibutyric urea (DSBU), which reacts with the side chains of lysine, serine, threonine and tyrosine residues. Crosslinked sites were then identified using MS and mapped onto the structure of TF, revealing the position of the NC relative to TF domains (Fig 2A and Table S4). TF consists of an N-terminal ribosome-binding domain (RBD) containing a conserved ribosome-binding site (RBS, _44_FRK_46_), a peptidyl-prolyl isomerase domain (PPD), and a C-terminal domain (CTD) (Fig 2B) ^13,56,57^. Most crosslinks between TF and the NC involved residues of the CTD, which was previously identified as the primary chaperone module in TF ^58^ (Fig 2C). The PPD also crosslinked promiscuously, to all RNCs except RNC_1-510_ and RNC_1-745_ which do not bind TF stably (Fig 1E). Crosslinks to the RBD were infrequent, and only observed in RNC_1-305_, RNC_1-352_ and RNC_1-687_.

**Figure 2.**
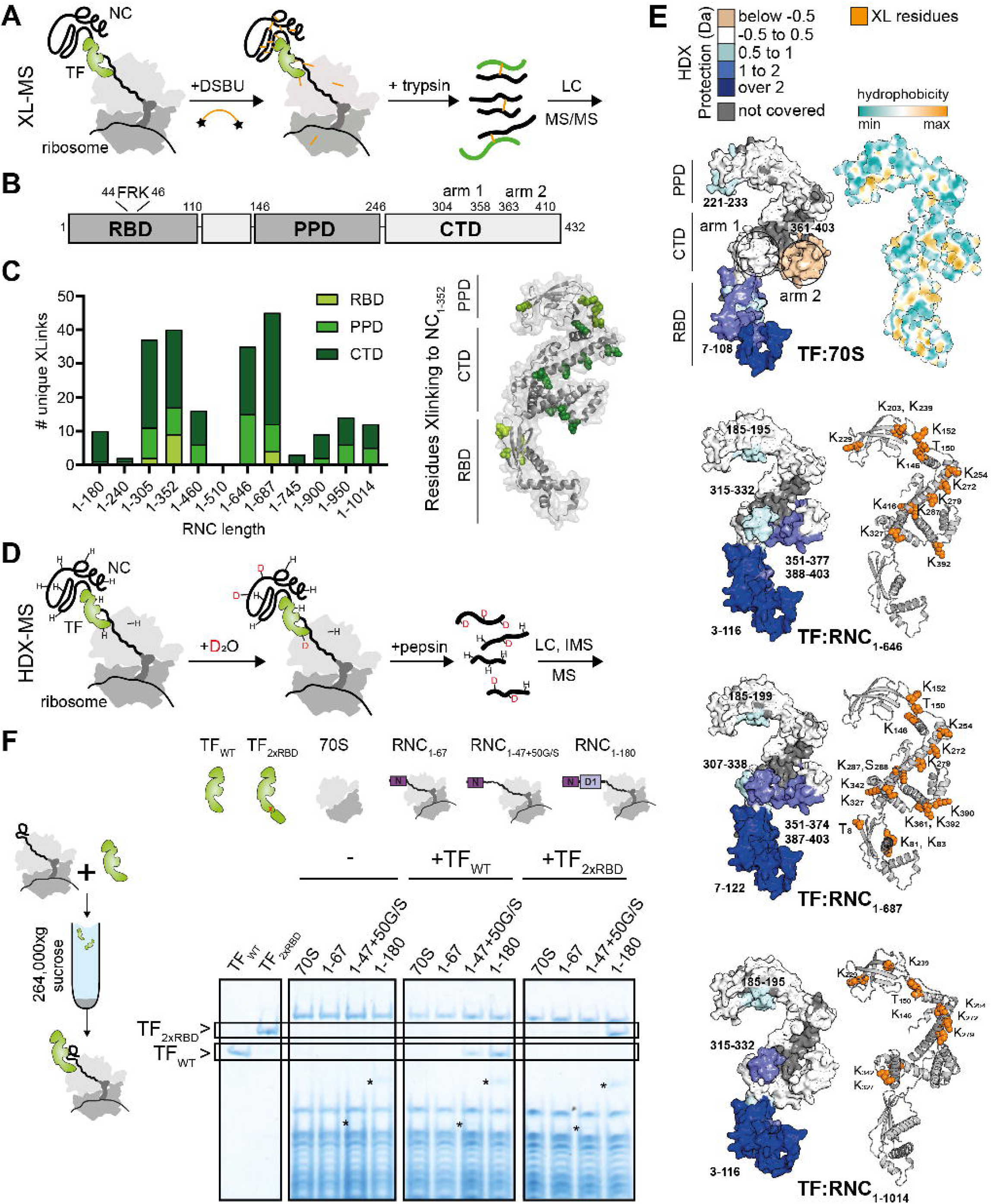
Nascent chains bind a versatile cavity created by the CTD and PPD of TF. **(A)** Schematic overview of the crosslinking-mass spectrometry (XL-MS) experiment. **(B)** Domain organisation of Trigger factor (TF) consisting of the ribosome-binding domain (RBD) with ribosome-binding site (_44_FRK_46_), peptidyl-prolyl isomerase domain (PPD) and the discontinuous C-terminal domain (CTD) containing the arm 1 and arm 2 regions. **(C)** Left: number of unique crosslinks detected between each TF domain and the NC of different β-gal RNCs (left). Right: residues on TF which crosslinked to NC_1-352_ are shown as green spheres on the TF monomer structure (PDB: 1W26 ^56^). **(D)** Schematic overview of the hydrogen/deuterium exchange-mass spectrometry (HDX-MS) experiment. **(E)** Left: TF monomer structures (PDB: 1W26) are coloured according to the difference in deuterium uptake (after 10 or 100 s deuteration) between isolated TF, and TF bound to 70S ribosomes, RNC_1-646_, RNC_1-687_ or RNC_1-1014_. Darker blue denotes less deuteration of bound versus isolated TF. Right: residues on TF that crosslink to each NC are shown as orange spheres. Top right: TF surface colour-coded according to hydrophobicity (plotted in ChimeraX ^91^). **(F)** Left: schematic overview of the TF:RNC co-sedimentation assay. Right: Coomassie-stained SDS-PAGE analysis of resuspended ribosomal pellets from co-sedimentation assays. Empty ribosomes (70S) or different β-gal RNCs (1-67, 1-47+50G/S, 1-180) were incubated with either wild-type TF (TF_WT_) or TF with two ribosome-binding domains (TF_2xRBD_). NCs are indicated (*) where visible. See also Fig S2 and Table S4,5

To validate the contribution of individual TF domains to β-gal RNC binding, we performed RNC co-sedimentation assays using purified TF mutants (Fig S2A). Deletion of the PPD prevented TF co-sedimentation with RNC_1-305_, RNC_1-352_ and RNC_1-900_ in high-but not low-salt buffer, indicating that the PPD contributes hydrophobic NC-binding sites (Fig S2B). The PPD was not required for TF binding to the four strongest interacting RNCs (RNC_1-180_, RNC_1-646_, RNC_1-687_ and RNC_1-1014_). Isolated (SUMO-tagged) TF RBD bound no more stably to RNC_1-352_ or RNC_1-687_ than to empty 70S ribosomes, indicating that RBD lacks high affinity NC binding sites (Fig S2C). Together with the XL-MS analysis, these data show that NCs primarily bind TF via the CTD and PPD, regardless of NC length.

To delineate the NC binding surface on TF more precisely, we measured RNC-induced protection of amide hydrogens in TF using hydrogen/deuterium exchange-mass spectrometry (HDX-MS) (Fig 2D). Compared to isolated TF, TF bound to RNC_1-646_, RNC_1-687_ or RNC_1-1014_ was protected from deuterium exchange in the RBD (Fig 2E and S2E and Table S5). We attribute this protection to ribosome rather than NC binding, since the same region was protected when TF was incubated with empty 70S ribosomes. Weak protection in 70S-bound TF was also observed at the tip of the PPD, suggesting that this site might form ribosome contacts in addition to the well characterised interaction between the TF RBD and ribosomal protein L23 ^13^.

Several other sites were uniquely protected in RNC-bound TF, and we attribute these to NC contacts (Fig 2E and S2E). All 3 NCs protected a hydrophobic patch on the inner surface of the PPD, supporting the idea that this domain contributes a universal hydrophobic NC binding site. In contrast, the pattern of protection in the CTD varied depending on the NC. TF bound to RNC_1-646_ was preferentially protected in the hydrophobic arm 2. In contrast, TF bound to RNC_1-1014_ was protected in the relatively hydrophilic arm 1, while TF bound to RNC_1-687_ was protected in both arms. The HDX data were consistent with XL-MS, as arm 1 crosslinked to all three RNCs, but arm 2 did not crosslink to RNC_1-1014_ (Fig 2E). Together, these data confirm that the CTD and PPD comprise the main NC binding surface, and reveal that the CTD contains multiple binding sites with mixed hydrophobic/hydrophilic character that differentially engage diverse NCs.

We hypothesised that the lack of stable binding to the ribosome-proximal RBD could allow this domain to act as a physical spacer, explaining a prior observation that TF is not recruited to NCs shorter than ∼100 residues *in vivo* ^7^. To test this, we measured TF binding to RNC_1-67_ and RNC_1-47+50G/S_ (Fig 2F). Both RNCs expose the same 47-residue N-terminal region of β-gal outside the exit tunnel, but RNC_1-47+50G/S_ contains an additional unstructured G/S-rich linker, which in an extended conformation is long enough to span the RBD. Note that the G/S linker alone does not stabilise TF binding, as neither RNC_40G/S_ nor RNC_FL+50G/S_ recruit TF (Fig 1E). We found that RNC_1-47+50G/S_ but not RNC_1-67_ bound TF, demonstrating that even a relatively short (47 aa) NC segment suffices to recruit TF, as long as the segment is physically extended beyond the RBD. A prediction of this model is that the size of the RBD is a critical determinant of the onset of TF binding. To test this prediction, we prepared a TF variant with a duplicated RBD (TF_2xRBD_), such that the distance between the exit tunnel and TF CTD is increased (Fig S2A,D). The second RBD in TF_2xRBD_ was mutated to prevent ribosome binding. TF_2xRBD_ bound RNC_1-180_ but not RNC_1-67_ or RNC_1-47+50G/S_, confirming that extending the RBD caused the onset of binding to shift to longer chain lengths (Fig 2F). Thus, the architecture of TF discriminates against short NCs due to their inability to reach the high-affinity binding cavity formed by the CTD/PPD.

### TF binds structurally compact nascent chains without antagonizing folding

Next, we analysed our XL-MS data to determine which parts of the NC are engaged by TF during synthesis. DSBU-reactive residues are distributed throughout β-gal, and we identified crosslinks to all 5 domains as well as the flexible N-terminal region (Fig 3A). Mapping these to each NC revealed that TF binding sites move from the N-terminal regions of the NC towards its C-terminus as translation proceeds (Fig 3B). This observation is consistent with TF remaining tethered to the ribosome exit port and suggests that the N-terminal domains in longer RNCs extend far enough to the cytosol to escape TF. Although the ribosome exit tunnel is expected to protect only ∼30 aa of the NC, just 3 of 12 NCs (RNC_1-352_, RNC_1-687_ and RNC_1-950_) crosslinked to TF via residues within 80 aa of the C-terminus (Fig S3A). This implies that, throughout translation, ribosome-proximal regions of the NC are disfavoured by TF, consistent with our conclusion that the RBD lacks high affinity NC binding sites.

**Figure 3.**
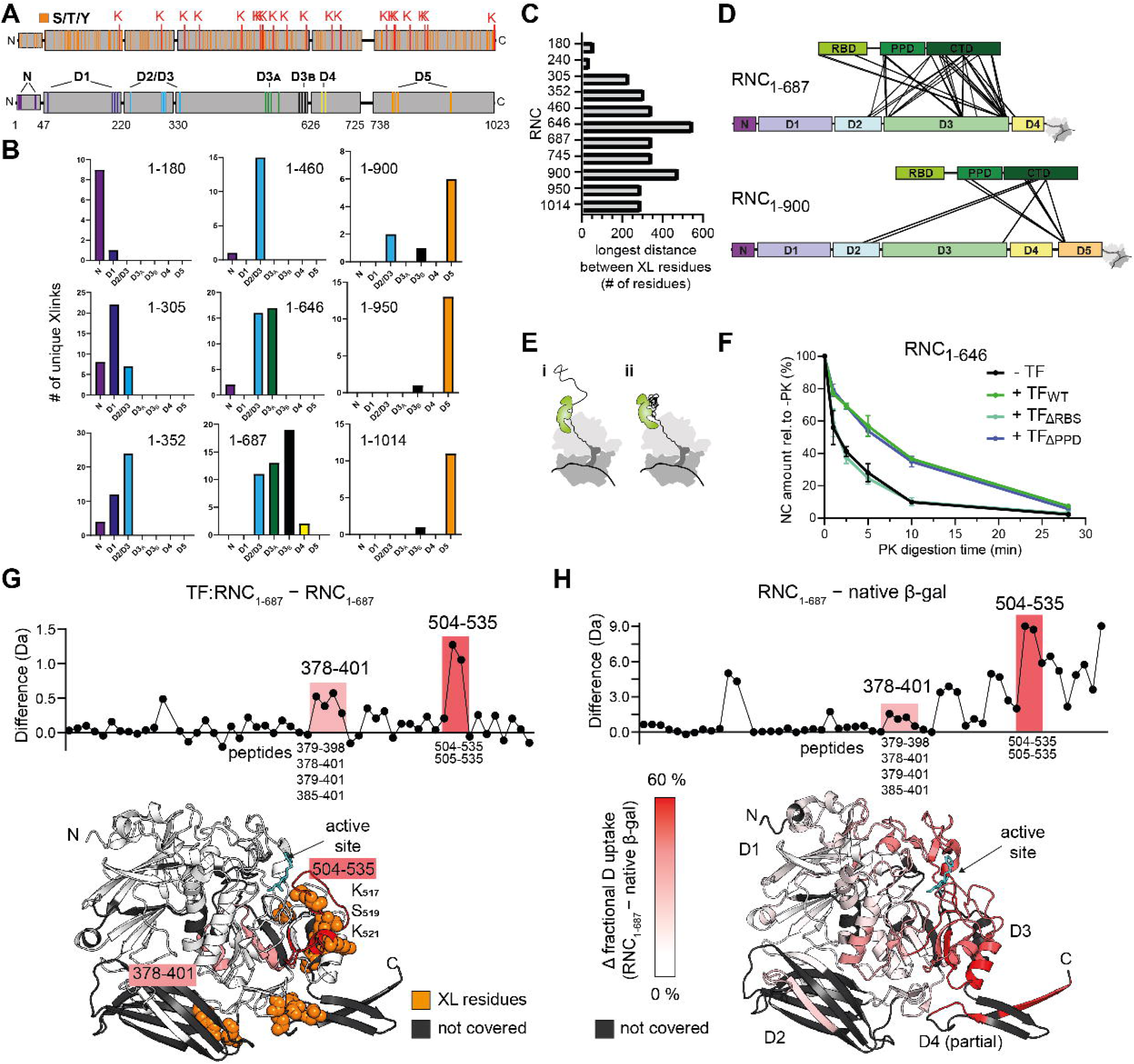
TF binds collapsed folding intermediates. **(A)** Top: β-gal domain organisation, highlighting DSBU-reactive residues. Lysine (K, red), serine, threonine, tyrosine (S, T, Y, orange). Bottom: color-coded grouping of residues in β-gal that crosslink to TF, based on their position relative to β-gal domains. **(B)** Number of unique crosslinks detected between TF and β-gal NC residues grouped as in (A). **(C)** Largest number of amino acids between two NC residues that crosslink to TF, for each RNC. **(D)** Crosslinks identified between TF and RNC_1-687_ or RNC_1-900_. **(E)** Possible TF binding modes. TF may bind NCs that are either highly extended (i), or collapsed and partially structured (ii). **(F)** Amount of undigested NC_1-646_ present at different time points after limited proteinase K digestion of RNC_1-646_ in isolation (-TF), or upon incubation with wild-type (TF_WT_) or mutated (TF_ΔRBS_ or TF_ΔPPD_) Trigger factor, based on SDS-PAGE analysis. Error-bars correspond to SD of triplicate reactions. **(G)** Top: difference in deuterium uptake, after 10 s deuteration, between RNC_1-687_ with and without bound TF. Values are plotted for individual peptides covering β-gal NC_1-687_, with regions 378-401 and 504-535 highlighted. Higher values indicate more deuteration in TF-bound RNC_1-687_ relative to free RNC_1-687_. Bottom: regions showing changes in deuterium uptake upon TF binding are mapped onto the structure of β-gal (PDB: 6CVM), truncated after 660 residues. Residues that crosslink to TF are shown as orange spheres. Regions that were not covered in the HDX-MS experiment are coloured dark grey. **(H)** Top: difference in deuterium uptake, after 10 s deuteration, between the NC in RNC_1-687_ and native β-gal. Higher values indicate more deuteration of RNC_1-687_ relative to native β-gal. Bottom: β-gal monomer structure as in (G), coloured according to the fractional deuterium uptake difference between RNC_1-687_ and native β-gal. Darker red indicates increased deuteration of peptides in RNC_1-687_ relative to native β-gal. The active site inhibitor PTQ is coloured cyan. See also Fig S3 and Table S4,5.

In all multidomain RNCs (RNC_1-305_ onwards) which stably bind TF, the CTD crosslinked to residues separated by >200 aa, from multiple domains, including residues far from the C-terminus (Fig 3C and S3B,C). This is exemplified by RNC_1-687_ and RNC_1-900_ which crosslinked to TF via 3 different β-gal domains, with crosslinks spread across 343 and 475 residues, respectively (Fig 3D and S3C). This crosslinking pattern could be explained by distant regions of the unfolded NC dynamically sampling the surface of ribosome-tethered TF, or indicate that the NC bound to TF is partially folded and highly compact (Fig 3E). To discriminate between these possibilities, we took several orthogonal approaches. First, we used HDX-MS to analyse RNC_1-646_ and RNC_1-687_ co-purifying with endogenous TF (Table S5). We found that 56-59% of the covered NC residues showed similar levels of deuterium exchange to native β-gal (<15% difference in fractional uptake), indicating that the NC is not substantially unfolded while bound by the chaperone (Fig S3D). Second, we analysed intra-β-gal crosslinks formed in TF bound RNCs (Table S4). These repeatedly formed between residues distant in sequence (up to 319 residues apart) but close together in native β-gal (Cα-Cα distance ≤ 35 Å ^59^) such as at the D2:D3 interface), consistent with the formation of native tertiary structure (Fig S3E). In the case of RNC_1-900_ exposing 4½ domains of β-gal, we also identified several crosslinks between D2/3 and D5, which are ∼70 Å apart in the native state. Incomplete D5 may thus occupy a non-native position during its synthesis such that overall chain is structurally compact while engaged by TF. Third, we tested the ability of TF to protect NCs from proteolytic cleavage. TF protected short (RNC_1-180_), medium (RNC_1-646_) and very long multidomain (RNC_1-1014_) RNCs from digestion by proteinase K (Fig 3F and S3F). Protection was dependent on the ability of TF to bind the ribosome, but did not require the PPD. Hydrophobic contacts between the NC and TF were not required, as the salt-sensitive binder RNC_1-460_ was similarly protected. Taken together, these observations are inconsistent with a model whereby TF interacts with extended, unfolded NCs. Instead, our data strongly suggest that NCs bound to TF are partially folded and structurally compact.

To determine the influence of TF on NC conformation, we purified RNC_1-646_ and RNC_1-687_ from cells lacking TF and analysed the complexes using HDX-MS (Table S5). Quantitative comparison with the equivalent TF-bound RNCs showed that NC deuterium uptake was globally unchanged by TF binding, although isolated regions were deprotected indicative of local conformational destabilization (Fig 3G and S3D,G). Residues 504-535 near the active site in D3 were strongly deprotected by TF binding in both RNCs, indicative of TF binding to this region, consistent with crosslinks between TF and residues 517, 519 and 521 of the NC. Residues 378-401 were weakly deprotected. Inspection of the mass spectra for peptide 504-535 revealed EX1 kinetics in the absence of TF, indicating that this region undergoes a concerted transition between low- and high-exchange states (Fig S3H). In the presence of TF, only the high-exchange state was detected, showing that TF selectively stabilises the less-folded conformation. Importantly, both regions affected by TF were already highly deprotected relative to native β-gal (Fig 3H). Thus, TF does not globally antagonize NC folding, but instead targets a subset of already non-native regions.

### TF is excluded from highly folded translation intermediates

To understand the determinants of TF binding to NCs, we focused on RNC_1-510_ (exposing 2½ domains) and RNC_1-745_ (exposing 4 domains) which do not stably bind TF (Fig 1E). RNC_1-510_ failed to recruit TF even upon introduction of a C-terminal linker to extend the NC further away from the ribosome surface, arguing against steric occlusion of the TF binding site on the ribosome (Fig S4A). The lack of TF binding was also not explained by competition with another cellular factor, as excess TF did not co-sediment with purified RNC_1-510_ or RNC_1-745_ *in vitro* (Fig 4A). Since the native state control RNC_FL+50G/S_ was not recognised by TF (Fig 1E), we hypothesized that stable folding of the incomplete NCs disfavoured TF binding. Indeed, introducing destabilising mutations ^60^ into the NC increased TF binding to both RNC_1-510_ and RNC_1-745_ (Fig 4A and S4B). This confirms that the binding mode of TF is influenced by the specific folding state of the NC, and suggests that partially synthesised intermediates with incomplete domains might adopt highly folded structures that exclude the chaperone.

**Figure 4.**
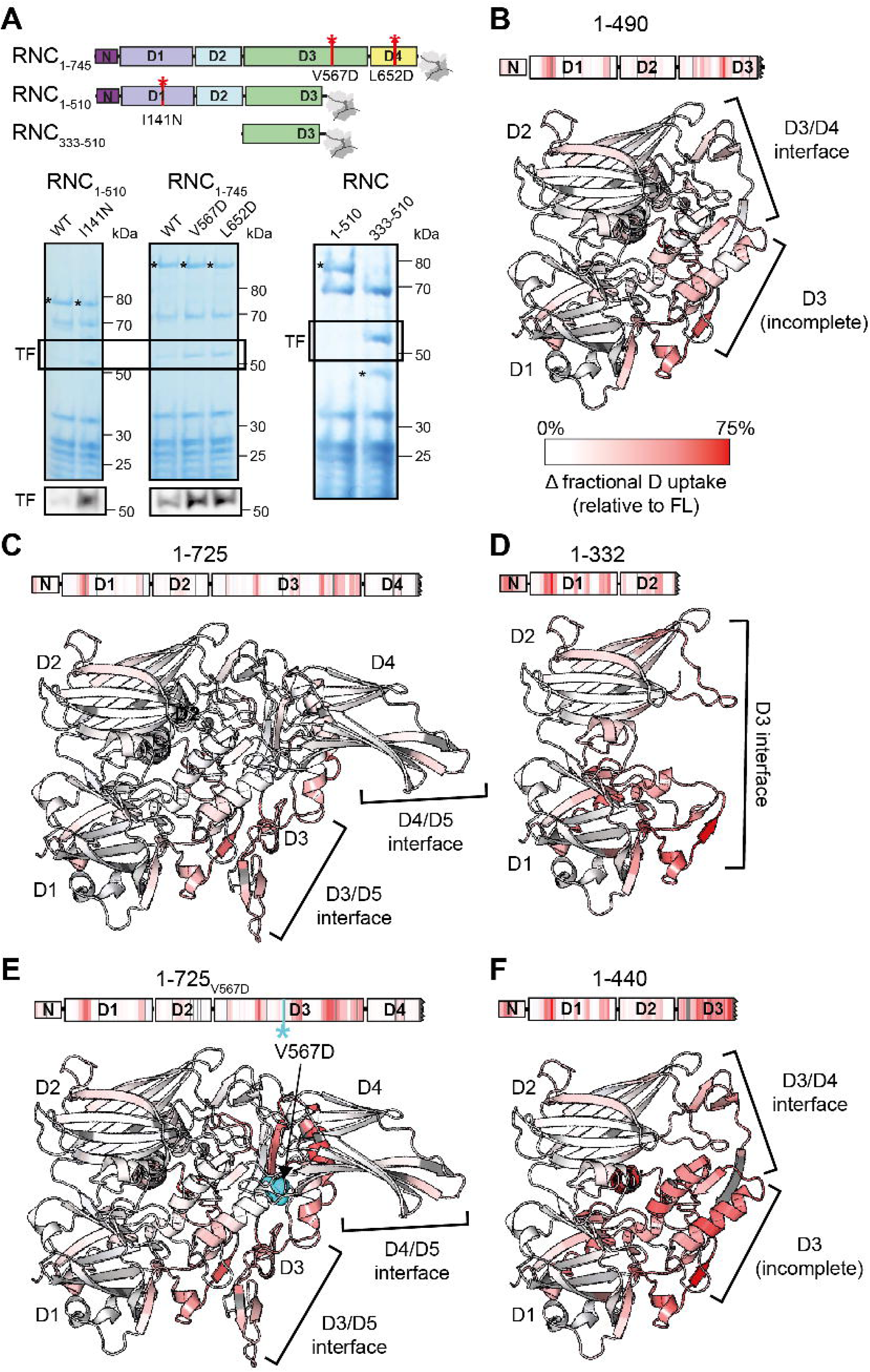
Structural determinants of TF binding to nascent β-gal. **(A)** Coomassie-stained SDS-PAGE gels of the resuspended ribosomal pellet from high-salt co-sedimentation assays of RNCs, purified from WT *E. coli* then incubated with additional TF *in vitro*. Bands corresponding to TF are highlighted. Bands corresponding to NCs (*) and TF are indicated. Immunoblots against TF are shown below. Positions of mutations are shown in red on the domain structure of NCs. **(B-F)** Fractional deuterium uptake difference, after 100 s deuteration, between full-length β-gal and β-gal chains truncated after 490 residues (B), 725 residues (C), 332 residues (D), 725 residues with an additional V567D mutation (E), or 440 residues (F). Darker red indicates more deuteration in the truncated chains compared to full-length β-gal. Orphaned domain interfaces are highlighted. Residue V567 is shown in cyan. See also Fig S4 and Table S6.

To characterise the conformation of TF-resistant and TF-binding NCs in detail, we took advantage of the fact that a subset of truncated chains, mimicking incomplete nascent polypeptides, were soluble in isolation (Fig S4C). We first analysed C-terminal truncations 1-490 and 1-725, which correspond to the part of the NC emerged from the exit tunnel in RNC_1-510_ and RNC_1-745_, respectively. HDX-MS showed that these chains adopt a near-native fold, with weak destabilisation (15-25% fractional uptake difference relative to native β-gal) at orphaned intra- or interdomain interfaces, and sites of strong destabilisation (>25%) clustered to adjacent loops in D3 and D1 around the active site (Fig 4B,C and S4D and Table S6). Thus, neither orphaned interfaces of near-native domains, nor localised non-native features on the NC surface suffice to recruit TF. For comparison we analysed 1-725_V567D_, 1-332 and 1-440, corresponding to NCs which do recruit TF (Fig 4D-F and S4D). In all cases, central elements in D1 or D3 were highly deprotected compared to native β-gal (>25% difference in fractional uptake), indicating incomplete domain folding.

Stable folding of C-terminal truncation 1-490 (corresponding to RNC_1-510_) was particularly striking, as the protein is truncated in the centre of the D3 TIM-barrel. To test whether an isolated half-barrel could escape TF, we prepared RNC_333-510_ containing only the N-terminal half of D3, without D1 or D2 (Fig 4A). Unlike RNC_1-510_, RNC_333-510_ strongly recruited TF, highlighting that TF engagement is determined not only by features of the newly-synthesized domain closest to the ribosome, but the complete NC including already-synthesized domains. Together, these data show that TF recognises partially folded (not necessarily partially synthesised) domains with extensive non-native features. Furthermore, domain-domain interactions can influence TF recruitment.

### DnaJ recruits DnaK to multiple sites on NCs

Other cytosolic chaperones besides TF are known to act at the ribosome ^41^. To capture less-stable interactors, we purified β-gal RNCs in low-salt buffer and omitted the second sucrose cushion centrifugation step. Under these conditions RNCs additionally co-purified with the chaperones DnaJ (Hsp40) and DnaK (Hsp70), both previously shown to engage NCs *in vivo* (Fig S5A)^9^. DnaJ and K were enriched over the negative control RNC_40G/S_ but present at low levels compared to TF (<10% occupancy), consistent with a more transient interaction (Fig S5B and Table S3). GroEL was also identified, but it was not substantially enriched over RNC_40G/S_.

To study the binding of DnaK and DnaJ to β-gal RNCs, we reconstituted the interactions *in vitro* using purified components (Fig 5A). DnaK alone did not co-sediment with RNC_1-646_ and required both DnaJ and ATP for stable binding. In contrast, DnaJ bound stably to RNC_1-646_, but was displaced by DnaK in the presence of ATP (Fig 5A and S5C and Table S7). Neither chaperone co-sedimented with empty 70S ribosomes under these conditions, confirming that the observed interactions depend on the NC (Fig S5D). The amount of co-sedimenting DnaK was reduced in the presence of low concentrations of the nucleotide exchange factor GrpE, and completely removed and accompanied by DnaJ rebinding when GrpE was added in large excess (Fig 5A and S5E). These results are consistent with the established model for client processing by the Hsp70 system ^61^ (Fig 5B). DnaJ stimulates binding of DnaK to substrate, dependent on ATP hydrolysis. GrpE then catalyses nucleotide exchange to release DnaK, allowing dynamic cycling of the chaperones *in vivo.* Although DnaJ and DnaK may be simultaneously accommodated on client proteins ^4,62–64^, our data suggest that the chaperones compete for binding to β-gal NCs. Except for RNC_FL+50G/S_ and RNC_1-67_, DnaK co-sedimented at a comparable level with every tested NC, including those that do not bind TF (Fig S5F-H). DnaK is therefore recruited to β-gal throughout synthesis but discriminates against natively folded and very short NCs.

**Figure 5.**
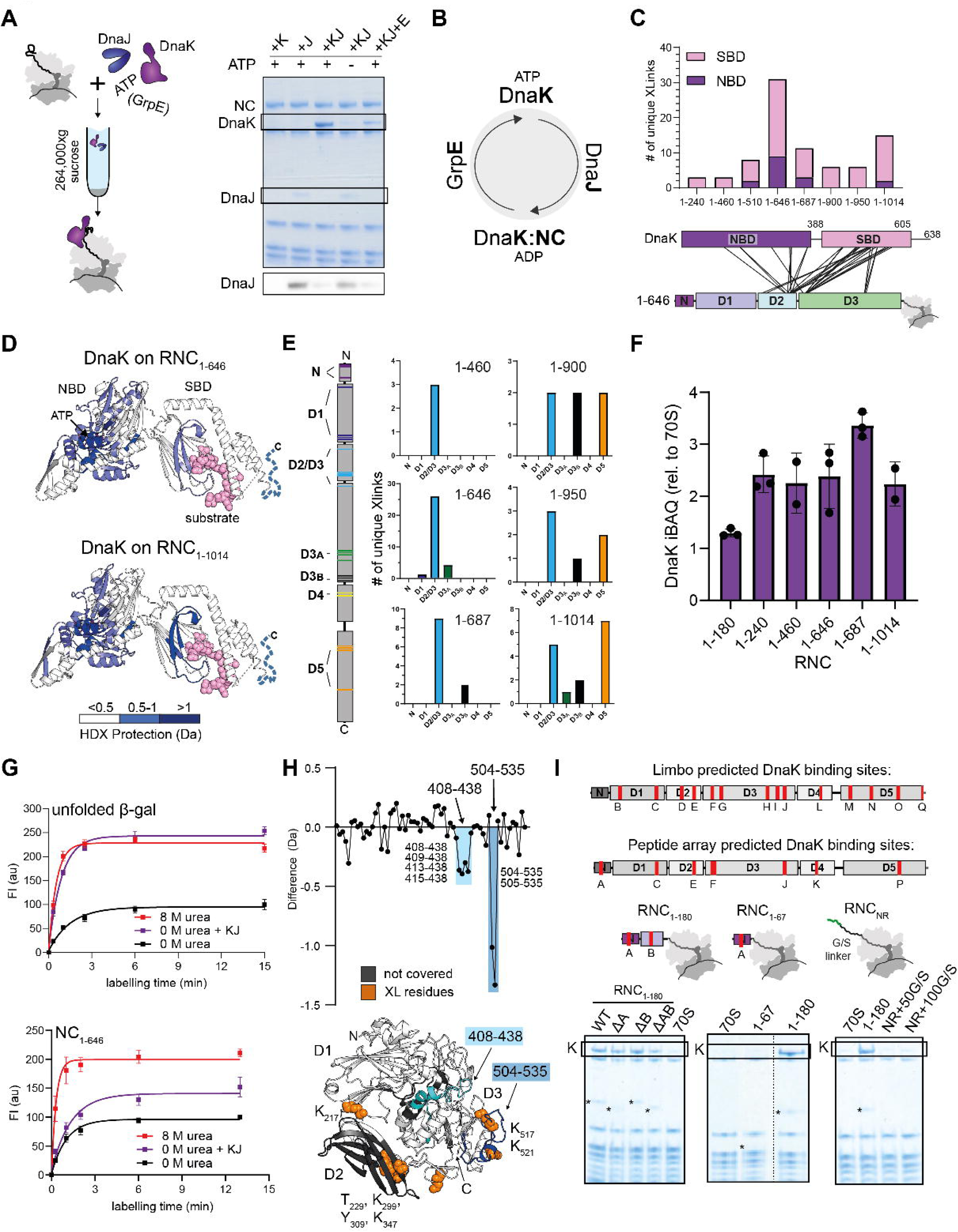
Architecture of complexes between DnaK and RNCs. **(A)** Left: schematic overview of the DnaJ/K:RNC co-sedimentation assay. Empty ribosomes or RNCs purified from ΔTF *E. coli* cells were incubated with DnaK, co-chaperones and ATP, then centrifuged through a 35% sucrose cushion to separate the ribosomal fraction (pellet) from unbound (co-)chaperones (supernatant). Right: Coomassie-stained SDS-PAGE of the resuspended ribosomal pellet. RNC_1-646_ was incubated with either DnaK (+K), DnaJ (+J), both DnaK and DnaJ (+KJ), or DnaK, DnaJ and GrpE (+KJ+E), with or without ATP as indicated. Bands corresponding to the NC_1-646_, DnaK and DnaJ are highlighted. An immunoblot against DnaJ is shown below. **_(B)_** Schematic of the canonical model for client processing by the Hsp70 system. DnaJ and GrpE control the transition of DnaK between conformations with low (ATP state) and high (ADP state) affinity for the client protein. Top: number of unique crosslinks between each DnaK domain (NBD - purple, SBD - pink) and the NC of different β-gal RNCs. Bottom: crosslinks identified between DnaK and RNC_1-646._ **(C)** Structure of peptide-bound DnaK (PDB: 7KZI ^92^) coloured according to the difference in deuterium uptake (after 10 or 100 s deuteration) between isolated nucleotide-free DnaK and ADP-bound DnaK in complex with RNC_1-646_ (top) or RNC_1-1014_ (bottom). Darker blue indicates less deuteration of RNC-bound DnaK relative to isolated DnaK. The C-terminal tail and linker not resolved in the structure are illustrated using dashed lines. The substrate peptide is shown as pink spheres. Bound ATP is shown in purple. **(D)** Number of unique crosslinks between DnaK and groups of residues on each NC. NC residues are grouped according to their position in β-gal domains, as in Figure 3A. **(E)** iBAQ values for DnaK present in the resuspended ribosomal pellet, following incubation of RNCs with DnaJ, ATP and excess DnaK. iBAQ values are normalised to the average iBAQ value of all 70S ribosomal proteins. Error-bars correspond to the SD of replicates from 2-3 independent co-sedimentation assays. **(F)** Cysteine-painting analysis of β-gal conformation. Samples containing isolated β-gal (top) or RNC_1-646_ (bottom) were incubated with Fluorescein-5-Maleimide for different times, resolved using SDS-PAGE, and the degree of β-gal labelling was quantified by fluorescent imaging. Prior to labelling, samples were incubated in buffer containing 0 M urea, 0 M urea with DnaK/DnaJ and ATP (0 M urea + KJ), or 8 M urea. Error-bars correspond to the SD of 3 independent labelling reactions. **(G)** Top: difference in deuterium uptake, after 10 s deuteration, between RNC_1-646_ with or without bound DnaK. Values are plotted for individual peptides covering NC_1-646_ with regions 408-438 and 504-535 highlighted. Lower values indicate less deuteration of peptides in DnaK-bound RNC_1-646_ relative to free RNC_1-646_. Bottom: β-gal monomer structure (PDB: 6CVM) truncated after 626 residues (corresponding to the NC region outside of the exit tunnel in RNC_1-646_) with regions 408-438 (cyan) and 504-535 (blue) highlighted. Residues on RNC_1-646_ that crosslink to DnaK are shown as orange spheres. Regions that were not covered in the HDX-MS experiment are coloured dark grey. **(H)** Top: position of candidate DnaK binding sites on β-gal (labelled A-Q) as determined by the LIMBO prediction algorithm ^67^ (Table S9) or an experimental peptide array. Bottom: Coomassie-stained SDS-PAGE of the resuspended ribosomal pellet from high-salt co-sedimentation assays of empty ribosomes (70S) or RNCs incubated with ATP, DnaJ and excess DnaK. RNC_1-180_ was mutated to remove site A (ΔA), site B (ΔB) or both sites (ΔAB). NR denotes the NR peptide sequence (NRLLLTG), positioned at the N-terminus of RNC_NR_ on a 50 or 100 residue G/S-rich linker. The band corresponding to DnaK is highlighted, and visible NCs are indicated (*). See also Fig S5 and Table S3,5,7-9.

To examine the molecular basis for DnaK recruitment to nascent β-gal, we first analysed DnaK:RNC complexes using XL-MS (Table S8) and HDX-MS (Table S5). The complexes were prepared by incubating RNCs with DnaK, DnaJ and ATP, followed by pelleting through a sucrose cushion. DnaK comprises a nucleotide binding domain (NBD) and substrate-binding domain (SBD), followed by an unstructured C-terminal tail. The vast majority of crosslinks between DnaK and NCs stemmed from the SBD (Fig 5C). Rare crosslinks to the NBD could be explained by interdomain flexibility in DnaK or originate from a second DnaK molecule bound to the same NC. HDX-MS analysis of DnaK bound to RNC_1-646_ and RNC_1-1014_ revealed protection, relative to isolated apo-DnaK, of the nucleotide binding pocket in the NBD and peptide binding groove in the SBD, as expected (Fig 5D and S6A). We also observed protection of the disordered C-terminal tail of DnaK (residues 601-630), previously shown to participate in substrate binding ^65^.

Sites on the NC that crosslinked to DnaK were not proximal in primary sequence or native structure, consistent with sparse binding of multiple molecules to the same NC (Fig 5C,E,H). Indeed, quantitative proteomic analysis showed that longer NCs co-sedimented with ∼2-3 molecules of DnaK on average (Fig 5F). Unlike TF, DnaK binding was not biased towards C-terminal parts of the NC, and N-terminal crosslinks were maintained long into translation (Fig 5E). Thus, DnaK persistently engages specific sites on the NC rather than simply binding newly-synthesized domains.

### DnaK does not stabilize an unfolded conformation of nascent **β**-gal

Simultaneous binding of multiple molecules of DnaK can stabilise unfolded conformations of substrate proteins ^31^. To probe the global conformation and tertiary structure of DnaK-bound NCs we used cysteine (Cys) painting, which covalently labels thiol groups with a fluorophore, dependent on their solvent exposure (Fig 5G) ^45^. Following labelling, reactions were separated by SDS-PAGE and subjected to fluorescence imaging, allowing the signal from β-gal to be distinguished from other proteins in the mixture. Full-length β-gal off the ribosome was labelled rapidly and extensively when denatured in 8 M urea but was relatively protected from Cys labelling following dilution from denaturant. Dilution from denaturant into a buffer supplemented with DnaK/DnaJ maintained β-gal in a globally unfolded state, indistinguishable by Cys painting from the urea-denatured state. In contrast, Cys labelling of β-gal NC in RNC_1-646_ was only slightly increased after incubation with DnaK/DnaJ/ATP. This was not due to competition for the reactive fluorophore by ribosomal proteins, as urea-denatured RNC_1-646_ was rapidly labelled. These data were complemented by peptide-level analysis using HDX-MS, which did not reveal any regions in NC_1-646_ that were deprotected upon DnaK binding (Fig 5H). Thus, despite binding stably to incompletely synthesised NCs, DnaK does not efficiently compete with NC folding. Since we focus on stable complexes, we do not exclude that NC folding is modified by transient interactions with DnaK. Nonetheless, our data establish that DnaK binding is compatible with NC folding.

### Canonical sequence motifs are neither necessary nor sufficient for stable DnaK binding to RNCs

Studies using isolated peptides have established that DnaK preferentially binds short (5-7 aa) predominantly hydrophobic sequence motifs ^66^. To systematically identify these motifs in β-gal, we both measured DnaK binding to a β-gal peptide array and annotated possible binding sites using the LIMBO server (Table S9) ^67^. This analysis resulted in a total of 17 potential binding sites, which we named A-Q (Fig 5I and S5I). To test the predictive quality of this approach for NCs, we focused on RNC_1-180_ which binds a single copy of DnaK on average (Fig 5F). Deleting the only peptide array-predicted site in RNC_1-180_ (site A, residues 1-21) did not affect the amount of co-sedimenting DnaK (Fig 5I). Similarly, DnaK binding was unaffected by mutating the only LIMBO-predicted site in RNC_1-180_ (site B, _57_EWRFAWF_63_ mutated to _57_EGSGSGF_63_). Combined mutation/deletion of both sites reduced, but did not eliminate, DnaK binding. Stable DnaK binding to NCs is therefore largely insensitive to the presence of canonical peptide motifs.

We next examined our HDX-MS data for evidence of DnaK binding, revealing two sites that were protected in the DnaK-bound state of NC_1-646_ (Fig 5H). The most prominent site was covered by residues 504-535 in D3, which includes residues K_517_ and K_521_ that crosslink to DnaK and contains a strong DnaK-binding motif as predicted by LIMBO (site H, residues _518_WSIKKWLSLP_527_, Fig 5I). Residues 408-426 in D3 were subtly protected by DnaK binding and might contain a second binding site, consistent with the expected 2:1 stoichiometry of the DnaK:NC_1-646_ complex (Fig 5F). Both sites mapped to loops that are solvent accessible in the structure of folded monomeric β-gal but buried at oligomeric interfaces in the native tetramer, which would explain why DnaK does not compete with monomer folding and is excluded from mature β-gal (Fig S5J). As observed for the predicted binding sites tested above, mutation of residues 504-535 (or just the LIMBO site 518-527) did not reduce the amount of DnaK co-sedimenting with RNC_1-646_ (Fig S5K). DnaK therefore binds NCs with little sequence specificity, and is apparently able to engage alternative sites depending on availability.

To probe the minimal requirements for stable binding, we additionally tested RNCs displaying the unstructured N-terminal extension of β-gal alone (RNC_1-67_, containing site A) or the “NR” sequence (NRLLLTG, a model DnaK-binding peptide^68,69^) appended to a 50- or 100-aa G/S-rich linker (Fig 5I). DnaK did not co-sediment with either RNC, indicating that a single DnaK-binding peptide motif does not suffice to engage and retain the chaperone during sucrose cushion centrifugation. Canonical short sequence motifs are therefore neither necessary nor individually sufficient for stable DnaK binding to RNCs, suggesting that additional factors regulate DnaK recruitment. Physical proximity of a J-domain has previously been shown to support promiscuous binding of Hsp70 to diverse peptides^70^. Since DnaK binding to RNCs is critically preceded by the J-domain protein DnaJ (Fig 5A,B), we next focused on DnaJ:RNC complexes.

### Architecture of complexes between RNCs and DnaJ

We first sought to identify the surface used by DnaJ (bacterial Hsp40) to bind RNCs. From N-to C-terminus, DnaJ consists of a J domain (JD) which stimulates ATP hydrolysis by DnaK, G/F-rich region (G/F), zinc binding domain (ZBD), two C-terminal β-sandwich domains (CTD I and II), and a dimerization domain (DD) (Fig 6A). Several Hsp40s have been shown to bind constitutively unfolded proteins via CTD I and II ^71^, although other domains have also been implicated in client recognition ^72–76^. It remains unclear how Hsp40s bind folding intermediates ^77^.

**Figure 6.**
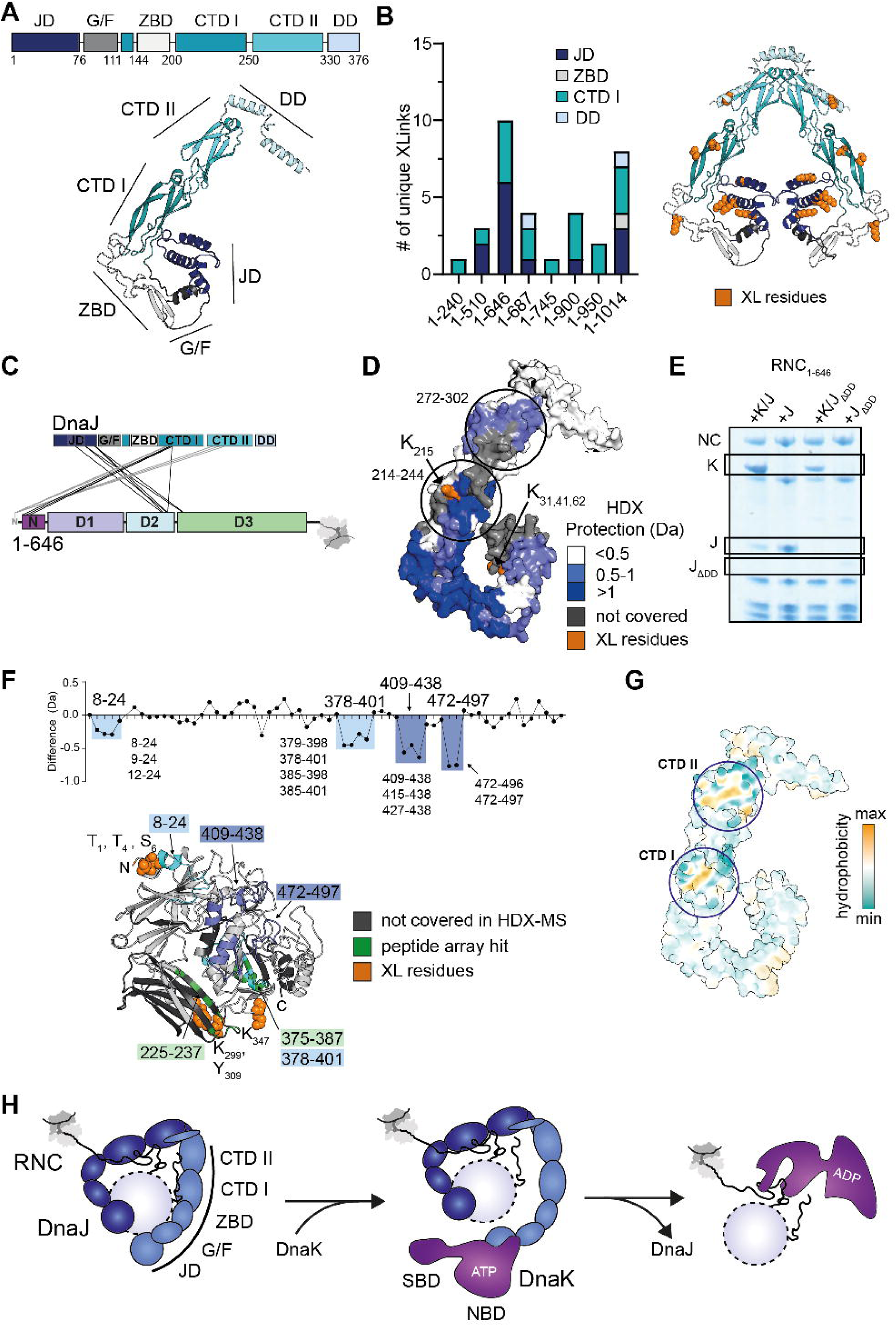
DnaJ binds RNCs using an extensive surface spread across multiple domains. **(A)** Domain organisation and predicted structure (AF-P08622-F1) of *E. coli* DnaJ monomer. The monomer consists of an N-terminal J domain (JD, dark blue), G/F-rich region (G/F, grey), zinc binding domain (ZBD, white), two C terminal β-sandwich domains (CTD I, aquamarine and CTD II, cyan), and a dimerization domain (DD, light blue). **(B)** Left: number of unique crosslinks between each DnaJ domain and the NC of different β-gal RNCs. Right: AlphaFold2.0 multimer-predicted structure ^93,94^ of dimeric DnaJ, coloured as in (A) and with residues that crosslink to any tested NC shown as orange spheres. **(C)** Crosslinks between DnaJ and RNC_1-646_. Crosslinks coloured in grey formed between DnaJ and the N-terminal amine of β-gal. Note that the 6 most N-terminal residues of purified RNCs (_1_GPGSGS_6_) encode a linker allowing efficient tag cleavage during purification. **(D)** Surface representation of DnaJ (AF-P08622-F1) coloured according to the difference in deuterium uptake (after 10 or 100 s deuteration) between isolated DnaJ and DnaJ bound to RNC_1-646._ Darker blue indicates less deuteration of RNC-bound DnaJ relative to isolated DnaJ. Residues on DnaJ that crosslink to NC_1-646_ are coloured orange. Regions that were not covered in the HDX-MS experiment are coloured dark grey. **(E)** Coomassie-stained SDS-PAGE of the resuspended ribosomal pellet from high-salt co-sedimentation assays of RNC_1-646_ incubated with either wild-type DnaJ (J) or DnaJ lacking the dimerization domain (J_ΔDD_), in the presence or absence of DnaK (K) and excess ATP. The bands corresponding to the NC, DnaK and DnaJ variants are highlighted. **(F)** Top: difference in deuterium uptake, after 100 s deuteration, between RNC_1-646_ with or without bound DnaJ. Values are plotted for individual peptides covering NC_1-646_ with regions 8-24, 378-401, 409-438 and 472-496 highlighted. Lower values indicate less deuteration of peptides in DnaJ-bound RNC_1-646_ relative to free RNC_1-646_. Bottom: β-gal monomer structure (PDB: 6CVM) truncated after 626 residues (corresponding to the NC region outside of the exit tunnel in RNC_1-646_) with protected regions coloured blue, sites detected in DnaJ peptide array coloured green, and residues that crosslink to DnaJ shown as orange spheres. Regions that were not covered in the HDX-MS experiment are coloured dark grey. **(G)** DnaJ monomer (AF-P08622-F1) surface colour-coded according to hydrophobicity (plotted in ChimeraX). **(H)** Model of DnaJ binding to RNCs and recruitment of DnaK. Dimeric DnaJ uses multiple domains from both monomers to bind NCs with mixed folded and unfolded character. During handover of substrates to DnaK, one DnaJ monomer may remain bound to the substrate, while the second monomer recruits DnaK via the DnaJ J-domain. DnaK binding to the NC displaces DnaJ See also Fig S6 and Table S4,5,8.

XL-MS analysis of DnaJ:RNC complexes identified frequent crosslinks to NCs via the JD and CTD I of DnaJ (Fig 6B,C and S6B). Crosslinks to CTD II were also observed, but exclusively to the extreme N-terminal amine of the NC (Table S8). The ZBD and DD formed a single crosslink with one and two RNCs, respectively. The G/F-rich region did not crosslink to any RNC, although we note that this region contains only one DSBU-reactive residue (Fig S6C).

We used HDX-MS as a complementary approach to map NC binding sites on DnaJ (Fig 6D and S6D, Table S5). Binding to RNC_1-646_ induced strong protection (>1 Da) in the G/F-rich region, ZBD and CTD I, all previously implicated in substrate binding ^71–76^. Weaker protection (0.5-1 Da) was observed in the JD, including the region containing the second most crosslinked residue K_62_, supporting its involvement in NC binding. A hydrophobic patch on CTD II (272-302) was also protected, but less so than the region protected in CTD I. The equivalent patch on *T. thermophilus* Hsp40 CTDII was previously shown to bind an unfolded polypeptide^71^) and is solvent accessible in dimeric DnaJ. Surprisingly, the strongly protected region we identify in CTD I (214-244, hereafter “site B”) is different to the previously-characterised peptide-binding site in the same domain (120-140, “site A”) ^71,78^ (Fig S6E). Our annotation of CTD I site B as a substrate binding site is supported by three additional observations. 1) Site B is next to the most crosslinked residue on DnaJ (K_215_, Fig 6D). 2) Site B forms a conserved hydrophobic groove structurally equivalent to the substrate binding site in the homologous CTD II (Fig S6F,G). 3) Alphafold 2.0 multimer predicts that site B is preferred by a model Hsp40-binding peptide GWLYEIS ^78^ (Li et al, 2003) (Fig S6E). Use of this alternative interaction site could be dictated by the overall topology of the binding surface, and might reflect differences in binding modes between constitutively unfolded proteins/peptides and compact folding intermediates. Recent work showed that peptides derived from p53 bind the equivalent site in yeast DnaJA2 (Ydj1), suggesting that site B is a conserved peptide-binding interface among class A J-proteins ^76^.

Neither XL-MS nor HDX-MS analysis pointed to a direct role for the DD in NC binding. However, the occurrence of homo-links (K_215_-K_215_ and K_31_-K_31_) in DnaJ bound to RNC_1-646_ suggested that DnaJ is bound to NCs as a dimer (Fig S6H). We found that removing the DD severely compromised both the binding of DnaJ to NCs and its ability to recruit DnaK, highlighting the functional importance of DnaJ dimerization (Shi et al, 2005) (Fig 6E).

We next sought to identify DnaJ-binding sites in nascent β-gal. HDX-MS revealed four sites on NC_1-646_ that were protected from deuterium uptake in the complex with DnaJ (Fig 6F), all of which were non-native in unbound NC_1-646_ (Table S5). No regions were deprotected by DnaJ binding. Weak but consistent protection occurred at residues 378-401 in D3, and the N-terminal region (8-24) to which CTD I formed multiple crosslinks (Fig 6C). Stronger protection occurred at residues 409-438 and 472-497 in D3. To complement these data, we measured DnaJ binding to a β-gal peptide array, which yielded two hits (Fig S6I). Peptide 375-387 overlapped with the HDX-protected region 378-401, while peptide 225-237 was not covered in the HDX-MS data but is close to several crosslinked residues, suggesting that both regions are genuine interaction sites in the intact protein (Fig 6F).

In summary, these data suggest that the JD, ZBD, G/F-rich region and hydrophobic groove on CTD I together comprise the primary NC binding surface on DnaJ, with CTD II potentially engaging the flexible N-terminus of the NC (Fig 6C,F,H). The resulting substrate-binding cradle has a mixed polar/hydrophobic character (Fig 6G) and, because of the unstructured G/F-region, is expected to be conformationally dynamic ^79^. These features may allow the chaperone to adapt to diverse partially-folded clients, and engage locally non-native sites on otherwise well-folded NCs. The fact that DnaJ dimerization is required for stable binding to NCs suggests that the binding interface spans both monomers. Thus, DnaJ might activate DnaK using the JD of one monomer while the other monomer engages the NC (Fig 6H). We show that the complex between monomeric DnaJ and the NC is relatively unstable, explaining why DnaJ is fully displaced upon DnaK recruitment (Fig 6E). Sites on the NC that were protected by DnaJ occurred nearby sites protected by DnaK, with only partial overlap (Fig S7A). This suggests that DnaJ and DnaK can bind neighbouring sites instead of directly competing for the same peptides, potentially facilitating substrate handover from DnaJ to DnaK.

### Coordination of TF and DnaK/J binding to RNCs

Since TF, DnaK and DnaJ all stably interact with nascent β-gal, we sought to investigate how these chaperones coordinate or compete at the ribosome. We noticed that upon TF knockout, some RNCs co-purified with increased amounts of DnaJ, pointing towards direct competition between TF and DnaJ for overlapping binding sites (Fig 7A). This phenomenon was replicated *in vitro,* where the addition of TF reduced the amount of DnaJ that co-sedimented with the same RNCs (Fig S7B). Three other observations support the idea that TF and DnaJ physically compete at the exit port. 1) Both TF and DnaJ (but not DnaK) repeatedly crosslinked to the solvent-exposed surface of ribosomal protein L29 at the exit port (Fig S7C and Table S4,8). 2) TF and DnaJ affect deuterium uptake in the same region of β-gal NCs (Fig S7A). 3) TF and DnaJ are similarly able to protect NCs from limited proteolysis (Fig 3F and Fig S7D).

**Figure 7:**
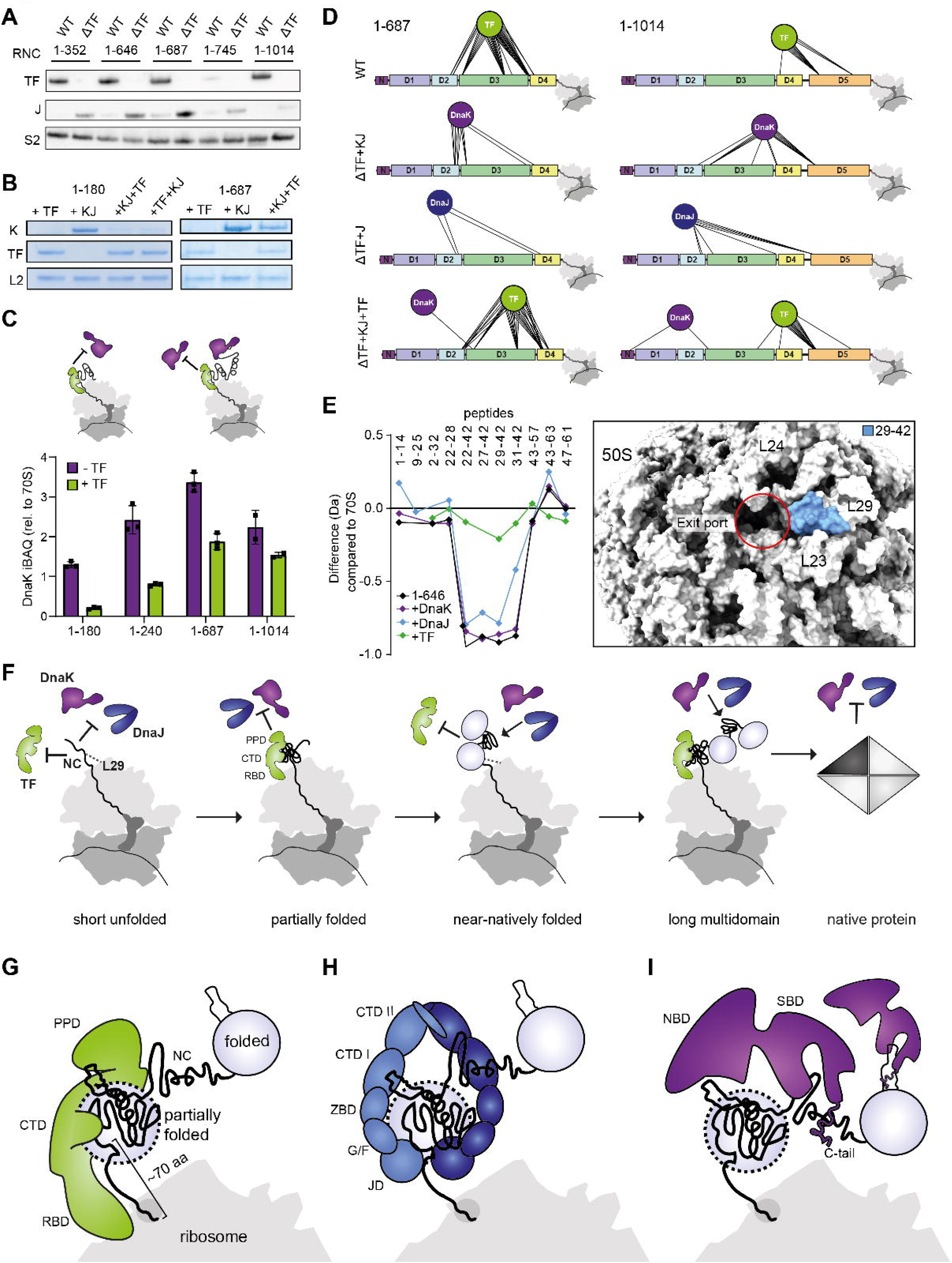
Coordination of TF, DnaJ and DnaK during multidomain protein synthesis. **(A)** Immunoblot of RNCs purified from either wild-type (WT) or ΔTF *E. coli* cells, probed for TF and DnaJ. Ribosomal protein S2 served as a loading control. **(B)** Coomassie-stained SDS-PAGE of the resuspended ribosomal pellet from high-salt co-sedimentation assays of RNC_1-180_ and RNC_1-687_, incubated with either TF (+TF), DnaK and DnaJ (+KJ) or all three factors in the specified order (+KJ+TF or +TF+KJ), all in the presence of ATP. Only the bands corresponding to DnaK, TF, and ribosomal protein L2 used as a loading control are shown. **(C)** iBAQ values for DnaK present in the resuspended ribosomal pellet from co-sedimentation assays of RNCs incubated with DnaJ, DnaK and ATP in the presence (green) or absence (purple) of TF. iBAQ values are normalised to the average iBAQ value of all 70S ribosomal proteins. Error-bars correspond to the SD of 3 independent co-sedimentation assays. The -TF data (purple bars) are the same as plotted in Fig5F. **(D)** Crosslinks between different chaperones and the NC in RNC_1-687_ (left) and RNC_1-1014_ (right). WT denotes RNCs purified from WT cells; ΔTF+KJ denotes RNCs purified from ΔTF cells then incubated with DnaK, DnaJ and ATP; ΔTF+J denotes RNCs purified from ΔTF cells then incubated with DnaJ; ΔTF+KJ+TF denotes RNCs purified from ΔTF cells then incubated with TF, DnaK, DnaJ and ATP. After crosslinking, samples were centrifuged through a high-salt sucrose cushion to remove unbound factors. **(E)** Left: difference in deuterium uptake, after 10 s deuteration, between isolated 70S ribosomes and either RNC_1-646_ (black) or chaperone-bound RNC_1-646_ (DnaK – purple, DnaJ – blue, TF – green). Values are plotted for individual peptides covering ribosomal protein L29. Lower values indicate less deuteration of L29 peptides in RNCs relative to isolated empty 70S ribosomes. Right: zoomed-in view of the ribosome exit port (red, PDB: 7D80 ^95^) with L29 residues 29-42 coloured blue. **(F)** Model of chaperone coordination at the ribosome during multidomain protein synthesis. Short NCs are not recognised by chaperones and can instead sample the ribosome surface (e.g. L29). TF is recruited when the NC elongates beyond ∼100 aa ^7^, allowing cotranslational folding in the CTD/PPD cavity and preventing DnaJ (and therefore DnaK) from binding. Near-natively folded intermediates exclude TF but are recognised by DnaJ, which then recruits DnaK to locally-unfolded sites. Longer NCs can simultaneously accommodate both TF and DnaK, resulting in persistent chaperone engagement of even NC segments that are far from the ribosome surface. Termination of synthesis and release from the ribosome allows the NC to escape TF, while domain docking and oligomeric assembly bury sites recognised by DnaJ/K. **(G)** Model of TF binding to RNCs. Collapsed, partially-folded NCs accumulate in the CTP/PPD cavity, exposing a mixed hydrophobic/hydrophilic surface. The binding cavity is physically separated from the exit tunnel by the TF RBD. **(H)** Model of DnaJ binding to RNCs. Unfolded regions bind hydrophobic grooves in the CTDs of DnaJ, while structured NC domains contact the predominantly hydrophilic ZBD, G/F-rich region and JD. The NC-binding interface spans both monomers of dimeric DnaJ. **(I)** Model of DnaK binding to RNCs. Solvent-exposed sites are bound by a narrow hydrophobic/hydrophilic groove in the DnaK SBD. The disordered C-terminal tail of DnaK also contacts the NC. See also Fig S7 and Tables S4,5,7,8.

We further studied the NC length-dependence of chaperone competition using *in vitro* pelleting assays. During synthesis of D1 (RNC_1-180_), TF outcompeted DnaK/J for NC binding, regardless of the order of addition or whether DnaK/J were added in excess (Fig 7B). Although the TF PPD contributes to RNC binding (Fig S2B), it was not required for TF to outcompete DnaK (Fig S7E). The ability of TF to suppress DnaK binding was progressively reduced during synthesis of D2 (RNC_1-240_, RNC_1-305_ and RNC_1-352_) even though TF levels were unaffected, suggesting that DnaK and TF can bind NCs simultaneously (Fig 7B and S5F). Indeed, two different populations of DnaK (sensitive and insensitive to TF binding) were observed on longer RNCs (Fig 7B,C and S5F and Table S7).

We next probed the topology of multi-chaperone:RNC complexes using XL-MS (Fig 7D). DnaJ or DnaK (in the absence of TF) crosslinked to both N- and C-terminal sites on the NC, while TF preferred C-terminal sites. When all 3 chaperones were added to RNCs together, the distribution of crosslinks between TF and the NC was unaffected. In contrast, DnaK was excluded from C-terminal sites but remained crosslinked to sites closer to the N-terminus. These data support a simple physical model for chaperone competition at the exit port. Ribosome-bound TF blocks DnaJ (and therefore DnaK) access to the C-terminal part of NCs, while N-terminal domains in longer NCs become sterically accessible to DnaJ/K once they escape the TF CTD (Fig 7C).

Despite the fact that the C-terminal part of the NC was consistently highly conformationally destabilised relative to native β-gal (Fig 3H and Table S5), we found no evidence by HDX-MS or XL-MS for chaperone binding to this region. Nor did chaperones stably bind the shortest tested RNC_1-67_ (Fig S5H). This could be explained by the availability of sequence motifs in the NC, or the position of binding sites on the chaperone in the case of TF (Fig 2F). Another possibility is that ribosome-proximal parts of the NC interact with the ribosome surface, competing with chaperone binding. To explore this idea, we analysed deuterium uptake in ribosomal proteins at the exit port, in the absence and presence of an NC and different chaperones (Fig 7E). We found that the exposed loop of ribosomal protein L29 was protected from deuterium exchange in RNC_1-646_ and RNC_1-687_ relative to empty ribosomes, consistent with a direct interaction between the NC and ribosome (Table S5). NC-induced protection of L29 was completely relieved by the binding of TF, suggesting that TF efficiently outcompetes the ribosome and routes the NC away from the ribosome surface. In contrast, neither DnaJ nor DnaK influenced L29 protection, indicating that these chaperones do not compete with the ribosome surface for NC binding.

## DISCUSSION

### Molecular basis for sequential chaperone action during *de novo* folding

Our data support a model in which the length and folding state of the emerging NC dictate the order and position of chaperone engagement (Fig 7F). At the earliest stages of translation (< 100 aa synthesised), the NC is not recognised by TF or DnaK/J, but can sample the ribosome surface near the exit tunnel. As the chain elongates, incomplete domains exposing substantial hydrophobic surface are stably bound by TF via multiple low-affinity sites. Burial of hydrophobic surface in an otherwise non-native intermediate does not release TF, which can also use electrostatic interactions to maintain loose contact with the NC. TF is only excluded by extensive NC folding, which can occur at intermediate points during synthesis. Remaining locally destabilized sites (e.g. interdomain regions or subunit interfaces) are recognised by DnaJ and then stably bound by DnaK, with a residence time controlled by GrpE-catalysed nucleotide exchange. Upon translation termination, the full protein sequence becomes available, allowing post-translational folding of C-terminal domains and completion of assembly interfaces, accompanied by exclusion of DnaJ/K.

The exclusion of chaperones at the earliest stages of translation could provide a defined temporal window of access for NC-modifying enzymes, as suggested previously ^7^. In addition, this delay would favour partial NC folding prior to chaperone engagement. Proximity to the bulky ribosome likely disfavours aggregation of short NCs, explaining why chaperones might not be required at this stage. Mechanistically, the delayed onset of TF binding is at least partially explained by the architecture of the chaperone, which allows it to function as a “molecular ruler” that measures the length of the NC (Fig 2). Short NCs may not satisfy the extensive interaction surface of the DnaJ dimer, explaining why neither DnaJ nor DnaK is recruited early in translation (Fig 6).

TF prevents binding of DnaJ/K to chains up to ∼200 residues (1 full domain of β-gal), consistent with previous observations that DnaK prefers NCs >30 kDa in size *in vivo*^15^. Thus, for smaller proteins, TF binding would enforce a purely post-translational engagement of DnaK. In longer, multidomain proteins, TF binding ensures that DnaJ/K can access only N-terminal domains that have escaped ribosome-bound TF (Fig 7). Priority binding of TF therefore increases the likelihood that DnaJ/K encounter substantially folded chains.

Since TF binding to diverse non-native folding intermediates can be driven by both hydrophobic and electrostatic interactions ^24,46^, a question remains as to how the chaperone discriminates between folded and unfolded NCs. One possibility is that TF binding is determined simply by the total solvent-accessible surface area of the NC, which reduces significantly upon folding ^80^.

### Architecture of the ribosomal exit port enforced by TF

We find that the TF RBD does not stably bind NCs, even though it contains a hydrophobic groove positioned directly adjacent to the ribosome exit port ^81,82^. We speculate that this groove weakly interacts with unfolded regions at the extreme C-terminus of the NC, while collapsed chains fold in the space between CTD/PPD and ribosome ^83^ (Fig 7G). This model is supported by our finding that the RBD crosslinked exclusively to residues near the C-terminus of NCs, while the CTD/PPD crosslinked to collapsed structures via sites hundreds of residues apart in sequence (Fig 3). In such a binding mode, TF effectively elongates the ribosome exit tunnel, keeping the C-terminal part of NCs in an extended conformation and insulating partially folded domains from destabilising contacts with the ribosomal surface ^40^ (Fig 7G). This could be especially significant for the maturation of multidomain proteins, which benefit from spatial separation of domains during folding ^5,19,84^. Interestingly, eukaryotic proteins tend to have much longer interdomain linkers than their bacterial counterparts ^85^, potentially explaining why TF is not conserved in the eukaryotic cytosol.

TF is highly abundant, has priority access to NCs, and is expected to engage most translating ribosomes ^7^. Thus, the exit port architecture enforced by TF is likely to be a general feature of cotranslational folding in bacteria.

### DnaJ and cotranslational folding dictate DnaK recruitment to NCs

Co-immobilization of a J-domain was previously shown to dramatically expand the range of peptide sequences that could be stably bound by the Hsp70 BiP ^70^. Consistent with this mechanism, we find that stable binding of DnaK to β-gal NCs is both highly promiscuous and depends on DnaJ. Rather than primary sequence bound by DnaK, DnaJ is the critical specificity factor for DnaK recruitment to NCs.

Client selection by Hsp40s is not well understood, but is generally assumed to rely on the availability of exposed hydrophobic stretches ^64,71,77^. NCs likely expose a combination of unfolded, hydrophobic segments and near-native structures, and our data suggest that DnaJ binds NCs via a surprisingly large surface is not exclusively hydrophobic (Fig 7H). This is consistent with the ability of DnaJ to recognise substrates in different folding states ^71,76^, (Fig 6) and the fact that Hsp40s generally bind intact proteins with much higher affinity than isolated peptides ^64,77^. The binding mode of DnaJ to NCs is strikingly similar to that of TF, also an ATP-independent chaperone, potentially providing a molecular explanation for the well-established functional redundancy of TF and DnaJ/K ^11,15,16^.

As folding is much faster than translation ^86^, cotranslational folding is expected to limit DnaK to loops or persistently unfolded regions of the NC (Fig 7I). This is consistent with our observation that DnaK binding stoichiometry does not increase with NC length, despite the increase in number of predicted binding sites (Fig 5). The significance of cotranslational folding to DnaK recruitment may be further reinforced *in vivo*, as DnaK engagement is delayed by TF and rapidly cycled by GrpE.

### Chaperone binding does not necessarily compete with cotranslational folding

Based on studies using isolated peptides, constitutively unfolded or chemically denatured clients, chaperone binding is typically understood to be driven by linear segments containing hydrophobic residues that are buried in the native state of substrate proteins ^87^. In this paradigm, chaperone binding competes with client folding, and in some cases can lead to further unfolding ^17,18,22,30,31,82,88^. More recently, both Hsp70 and TF have been suggested to also bind in alternative modes which stabilise rather than unfold _clients_ 20,21,24,26.

We show that, during cotranslational folding, chaperones target partially folded nascent chains without reversing incipient folding. Evidence for this comes from multiple observations. First, HDX-MS analysis of NCs without bound chaperones reveals extensive native-like folding. This does not globally change upon TF, DnaJ or DnaK binding, and any local effects map to already-unfolded regions. Second, intra-NC crosslinks consistent with native tertiary structure do not change upon chaperone binding (Fig S7F). Third, both TF and DnaJ protect NCs from proteolytic degradation. Fourth, DnaK binding does not globally expose buried cysteines. Beyond preventing aggregation and non-specific degradation, continuous chaperone engagement could function to prevent inter-domain misfolding during synthesis.

### Limitations of the study

Although the structural proteomics approaches used in our study facilitate detailed analysis of large complexes with dynamic elements, they have several limitations. First, XL-MS does not distinguish between rare and frequently sampled conformations, a consideration that is especially pertinent for the structurally heterogenous NC. Second, the size and RNA-rich character of RNCs is a substantial challenge HDX-MS, limiting sensitivity and spatial resolution. Third, allosteric effects can complicate the identification of binding sites using HDX-MS, necessitating that the data are interpreted together with XL-MS and the results from binding experiments using mutant chaperones.

More generally, it remains to be determined whether protein maturation *in vivo* is faithfully recapitulated by isolated RNC:chaperone complexes, removed from the context of ongoing translation and transient binding of chaperones or NC-modifying enzymes. Other chaperones not studied here, such as GroEL/ES, may also modify NC maturation. Furthermore, the generality of our conclusions remains to be tested using additional NC models.

## METHODS

### DNA vectors and cloning

The DNA encoding full-length β-galactosidase was amplified from NEB *Escherichia coli* (*E. coli*) BL21(DE3) genome using the PureLink Genomic DNA Mini Kit (ThermoScientific) and cloned into a pET28 vector following an N-terminal His_6_-tag. Gibson Assembly cloning (NEB) was used to generate a set of pET21 plasmids encoding β-galactosidase ORFs upstream of a ribosome stalling sequence (WWPRIRGPPGS ^90^) and downstream of HRV 3C-cleavable monomeric ultrastable GFP (muGFP) tag ^105^. *E. coli* DnaK and DnaJ were expressed without any tags from pET11d, and GrpE from pET3a. His_6_-tagged TF was expressed from plasmid ProEX. Wild-type chaperone expression vectors were kind gifts from F.U. Hartl (MPI Biochemistry). The plasmid encoding a His_6_-SUMO-tagged TF variant with two ribosome-binding domains (TF_2xRBD_) was synthesised by Twist Biosciences and cloned into a pET28 backbone. Additional insertions, deletions and point mutations were introduced using site-directed mutagenesis with Q5 or Phusion polymerases (NEB). All constructs used in this study (Table S10) were verified by sequencing.

### RNC buffers

RNC low-salt buffer contained 50 mM HEPES-NaOH pH 7.5, 12 mM Mg(OAc)_2_, 100 mM KOAc, 1 mM DTT and 8 U/mL RiboLock RNase inhibitor (ThermoScientific). RNC high-salt sucrose cushion contained 35% sucrose, 50 mM HEPES-NaOH pH 7.5, 12 mM Mg(OAc)_2_, 1 M KOAc, 1 mM DTT, 8 U/mL RiboLock RNase inhibitor and 0.2x Halt Protease Inhibitor Cocktail (ThermoScientific). RNC low-salt sucrose cushion contained 35% sucrose, 50 mM HEPES-NaOH pH 7.5, 12 mM Mg(OAc)_2_, 100 mM KOAc, 1 mM DTT, 8 U/mL RiboLock RNase inhibitor and 0.2x Halt Protease Inhibitor Cocktail.

### Protein purification

#### Expression and purification of ribosome:nascent chain complexes (RNCs)

RNCs were expressed and purified as described previously ^46^ with slight modifications. In short, BL21(DE3) *E. coli* cells (NEB) were transformed with plasmids encoding β-gal RNC constructs and grown in ZYM-5052 autoinduction media ^106^ for 18 h at 37 °C. Cultures were pelleted (4,000 g, 30 min) and resuspended in RNC lysis buffer (70 mM Tris, pH 7.5, 150 mM KCl, 10 mM MgCl_2_, 8 U/mL RiboLock RNase inhibitor, 0.5x Halt Protease Inhibitor Cocktail, 2 mg/mL lysozyme, 0.05 Kunitz units/µL RNase-free DNase (QIAGEN)). Following an incubation (30 min, 4 °C) and subjecting resuspended pellets to at least 2 freeze-thaw cycles at -80 °C, the soluble fraction was separated by centrifugation (20 min, 16,000 g) and centrifuged (2 h, 264,000 g) through RNC high-salt sucrose cushion to isolate ribosomes in the pellet. The pellet was resuspended in RNC low-salt buffer and applied to in-house prepared GFP-clamp-agarose beads ^107^ for 16 hours at 4 °C. RNCs were selectively eluted by muGFP-tag cleavage with HRV 3C protease, and further purified by pelleting through a second sucrose cushion (2 h, 264,000 g). Pellets were resuspended in RNC low-salt buffer, snap-frozen in liquid nitrogen and stored at −80 °C. Sequences of purified proteins are listed in Table S11.

#### Expression and purification of isolated ***β***-galactosidase variants

BL21(DE3) *E. coli* Δ*lac* cells (Addgene, Didovyk et al., 2017) transformed with plasmids encoding β-gal (wild-type or mutant) were grown in LB at 37 °C until OD_600_ 0.6–0.8 and expression was subsequently induced with 1 mM IPTG for an additional 18 hours at 16-18 °C. Cells were harvested (4,000 g, 30 min), resuspended in Buffer A (50 mM Tris-HCl, pH 7.5, 500 mM NaCl, 10% glycerol, 2 mM β-mercaptoethanol, 100 mM PMSF) supplemented with 1 mg/mL lysozyme, benzonase (Millipore) and Complete EDTA-free protease inhibitor cocktail (Roche), and lysed by sonication. Lysate was clarified by centrifugation (60,000 g, 45 min) and then applied to HisTrap HP column (Cytiva) equilibrated in Buffer A. Peak fractions from gradient elution with 500 mM Imidazole were treated with TEV protease (48 h, 4 °C) and dialysed against Buffer A. Uncleaved protein was removed using a HisTrap HP column equilibrated in Buffer A. The flow-through was further purified using a Superdex 200i 10/300 or HiLoad Superdex 200 16/600 (Cytiva) equilibrated in Buffer B (20 mM Tris-HCl, pH 7.5, 300 mM NaCl, 10 mM MgCl_2_, 3 mM DTT, 5% glycerol), and pure protein was concentrated and snap-frozen in liquid nitrogen for storage at −80 °C. Sequences of purified proteins are listed in Table S11. Destabilising mutations of β-galactosidase, I141N and V567D, were previously characterised ^60^. Destabilising mutation L652D was designed using DynaMut2 ^102^.

#### Expression and purification of DnaK, DnaJ and GrpE

BL21(DE3) *E. coli* transformed with DnaK-, DnaJ- or GrpE-encoding plasmids were grown in LB at 37 °C until OD_600_ 0.6–0.8 and subsequently induced for expression with 0.5 mM IPTG for an additional 5 hours at 30 °C.

DnaK-expressing cells were harvested by centrifugation (4,000 g, 30 min), resuspended in lysis buffer (20 mM Tris-HCl, pH 7.4, 1 mM EDTA, 1 mg/mL lysozyme, 0.2 mM PMSF, benzonase, and Complete EDTA-free protease inhibitor cocktail) and lysed by sonication. Clarified lysate (60,000 g, 45 min) was applied onto RESOURCE Q column (Cytiva) preequilibrated in Buffer Q (20 mM Tris-HCl, pH 7.4, 1 mM EDTA). Peak fractions eluted with NaCl were combined and desalted on HiPrep 26/10 desalting column (Cytiva) equilibrated in Buffer H (20 mM HEPES-NaOH, pH 7.4, 5 mM Mg(OAc)_2_) and loaded onto a HiTrap Heparin column (Cytiva) equilibrated in Buffer H. Peak fractions eluted with an NaCl gradient were exchanged into Buffer Q and loaded onto a RESOURCE Q column. DnaK-containing fractions from NaCl elution were combined and loaded onto a Superdex 200i 10/300 column in 20 mM HEPES-NaOH, pH 7.4, 5 mM Mg(OAc)_2_, 100 mM NaCl, 5% glycerol, 1 mM DTT. Pure DnaK was concentrated, snap-frozen in liquid nitrogen and stored at −80 °C.

DnaJ-expressing cells were harvested by centrifugation (4,000 g, 30 min), resuspended in lysis buffer (50 mM Tris-HCl, pH 8.5, 10% sucrose, 10 mM DTT, 10 mM EDTA, 0.6% Brij58, 1 mg/mL lysozyme, benzonase, and Complete EDTA-free protease inhibitor cocktail) and lysed by sonication. The lysate was centrifuged (60,000 g, 45 min) and the pellet was resuspended in Buffer U (2 M urea, 50 mM Tris-HCl, pH 8.5, 5 mM DTT, 10% sucrose, 0.1% Triton-X). Resolubilised protein was separated from aggregated material in the pellet by centrifugation (60,000 g, 45 min), and the supernatant was loaded onto RESOURCE Q column in Buffer U. Peak fractions eluted with Buffer Q2 (50 mM Tris-HCl, pH 8.5, 5 mM DTT, 10% glycerol, 0.05% Brij58, 1 M NaCl) were combined and desalted using a HiPrep 26/10 desalting column equilibrated in Buffer S (20 mM sodium phosphate, pH 6.8, 5 mM DTT, 10% glycerol, 0.05% Brij, 100 mM KCl). Protein was subsequently loaded onto RESOURCE S column (Cytiva) preequilibrated in Buffer S. Protein eluted with KCl was exchanged into Buffer S, loaded onto a HiTrap Heparin column, and eluted with a KCl gradient. DnaJ-containing fractions were combined and loaded onto Superdex 200i 10/300 column in 25 mM Tris-HCl, pH 7.5, 10% glycerol, 5 mM DTT, 100 mM KCl. Pure DnaJ was concentrated and snap-frozen in liquid nitrogen for storage at −80 °C. The DnaJ mutant without the dimerization domain (DnaJ_ΔDD_ encoding residues 1-330) was purified following the same protocol.

GrpE was purified as described in ^30^ with the following modifications. Spermidine was omitted from the lysis buffer and a RESOURCE Q column was used for all anion-exchange steps. A Superdex 200i 10/300 column in Buffer E (25 mM HEPES-NaOH pH 8.0, 1 mM EDTA, 10 mM β-mercaptoethanol, 20% glycerol) was used for a size-exclusion chromatography step. Sequences of purified DnaK, DnaJ and GrpE proteins are listed in Table S11.

#### Expression and purification of TF

Trigger factor and its mutants (TF_ΔPPD_, TF_ΔRBS_, TF_2xRBD_, RBD_N-tag_, TF_R14C_ and TF_ΔRBS-R14C_, Figure S2A) were purified as described in ^46^ with slight modifications. In short, BL21(DE3) *E. coli* transformed with a plasmid encoding the appropriate TF variant were grown in ZYM-5052 auto-induction media ^106^ (18 h, 37 °C). Cells were harvested (4,000 g, 30 min), resuspended in lysis buffer (50 mM Tris-HCl, pH 7.5, 250 mM NaCl, 20 mM Imidazole, 1 mM DTT, 1 mg/mL lysozyme, benzonase and Complete EDTA-free protease inhibitor cocktail) and lysed by sonication. Clarified lysate (60,000 g, 45 min) was applied to HisTrap HP column equilibrated in Buffer T (20 mM Tris-HCl, pH 7.5, 1 mM DTT, 250 mM NaCl, 20 mM Imidazole). Peak fractions from gradient elution with 500 mM Imidazole were treated with TEV protease (48 h, 4 °C) and dialysed against Buffer T. Uncleaved TF was removed by reapplying protein onto HisTrap HP column equilibrated in Buffer T. Following further purification on a Superdex 200i 10/300 in size-exclusion buffer (20 mM Tris-HCl, pH 7.5, 150 mM NaCl, 1 mM DTT, 5% glycerol), pure protein was concentrated and snap-frozen in liquid nitrogen for storage at −80 °C. The TEV-cleavage step was omitted for RBD_N-tag_. The TEV-cleavage step was replaced with a Ulp1-cleavage step for TF_2xRBD_. Sequences of purified proteins are listed in Table S11. Mutations introduced in TF_ΔRBS_ (F44A-R45A-K46A), TF_R14C_ and TF_ΔPPD_ (replacement of residues 151-243 with GTSAAAG) were described previously ^13,51,52^.

### Biochemical assays

#### *β*-galactosidase activity assay

β-gal activity was measured in RNC low-salt buffer with 2.1 mM o-nitrophenyl-β-D-galactopyranoside (ThermoScientific) by following absorbance at 420 nm at 25 °C and recording the slope of the progress curve. Concentration of β-gal and RNCs in stock samples was 100 nM. Stock solutions were diluted 20-fold into assay buffer just before activity measurement. Activity of each stock sample was recorded in three independent measurements.

#### RNC:chaperone binding assays

For co-sedimentation binding assays, 2 µM RNCs or empty 70S ribosomes (NEB) were incubated (30 min, 30 °C) with (co-)chaperones (DnaK – 10 µM, TF – 10 µM, DnaJ – 2-5 µM) in RNC low-salt buffer supplemented with 5 mM ATP unless otherwise indicated. Samples were loaded onto a 35% high- or low-salt sucrose cushion as indicated, and pelleted by centrifugation (264,000 g, 2 h). Following two washes with ice-cold RNC low-salt buffer, the pellet was resuspended at 4 °C. Resuspended pellets from co-sedimentations assays were analysed by SDS-PAGE or proteomics analysis and where indicated used as input for other assays.

For fluorescence-based binding assays, TF_R14C_ and TF_ΔRBS-R14C_ were labelled with a cysteine-reactive fluorophore. Purified TF cysteine-mutants diluted to 100 µM in 1 mL of PBS were first reduced by TCEP (1 mM, 30 min, 25 °C) and subsequently labelled by incubating (90 min, 25 °C) with 5-fold molar excess of BADAN (Santa Cruz Biotechnology). The labelling reaction was quenched by addition of 10 mM DTT. After centrifugation (15,000 g, 20 min, 4 °C), excess reagent was removed by gel-filtration using PD-10 columns (Cytiva) following manufacturer instructions. The labelling efficiency was determined to be 70-80% by measuring absorbance at 387 nm (dye peak absorbance) and 280 nm (protein peak absorbance) and applying the appropriate correction to account for absorbance of BADAN dye at 280 nm. Labelled TF variants in assay buffer (10 mM Tris, pH 7.5, 10 mM MgCl_2_, 6 mM β-mercaptoethanol, 1 mg/mL BSA, 0.01% Tween) were used to detect the change in fluorescent intensity (λ_ex_ 384 nm, λ_em_ 508 nm) upon mixing with RNC_1-646_, as previously described ^51^.

#### Proteinase K assay

RNCs at 0.5 µM were incubated with 1 µM of TF or 2 µM DnaJ dimer in RNC low-salt buffer for 30 min at 30 °C, then cooled to 4 °C. A reference aliquot (t_0_) was removed before initiating degradation at 4 °C by addition of Proteinase K (Millipore) to a final concentration of 2.5 ng/µL. Aliquots of reactions were removed and quenched at different times by 1:1 mixing with 5 mM PMSF in RNC low-salt buffer, then analysed by SDS-PAGE and Coomassie staining. Quantification of the tRNA-NC band was performed in Fiji ^97^ and values were plotted as a mean of three replicate reactions.

#### Cysteine painting assay

Initially, all proteins were buffer exchanged into assay buffer (50 mM HEPES-NaOH pH 7.5, 12 mM Mg(OAc)_2_, 100 mM KOAc, 8 U/mL RiboLock RNase inhibitor, 5 mM ATP) using Micro Bio-Spin 6 columns (BioRad) according to the manufacturer’s instructions. To prepare unfolded samples, β-galactosidase and RNC_1-646_ were incubated (4 h, 25 °C) in assay buffer supplemented with 8 M urea. Unfolded β-galactosidase (0.5 µM) was subsequently incubated (30 min, 30 °C) in assay buffer, assay buffer with DnaK (5 µM) and DnaJ (2.5 µM), or assay buffer with 8 M urea. Additionally, untreated RNC_1-646_ (0.5 µM) was incubated (30 min, 30 °C) in assay buffer with or without DnaK (5 µM) and DnaJ (2.5 µM). Following the above-described incubations, cysteine painting of β-galactosidase and RNC_1-646_ samples was initiated at 10 °C by rapid mixing with Fluorescein-5-Maleimide (ThermoScientific) at a final concentration of 0.1 mM. Aliquots from triplicate labelling reactions were removed at different time points and quenched by mixing with excess SDS-PAGE loading dye containing β-mercaptoethanol. Quenched reactions were resolved on SDS-PAGE gels and Fiji ^97^ was used for quantification of the fluorescence intensity and Coomassie stain intensity of specific bands. Fluorescence intensity was normalised to Coomassie stain intensity to account for loading errors.

#### Peptide arrays

Peptide arrays were synthesised on an Intavis ResPepSL Automated Peptide Synthesiser (Intavis Bioanalytical Instruments) on a cellulose membrane by cycles of N(a)-Fmoc amino acids coupling via activation of carboxylic acid groups with diisopropylcarbodiimide (DIC) in the presence of Ethyl cyano(hydroxyimino)acetate (Oxyma pure) followed by removal of the temporary α-amino protecting group by piperidine treatment. After chain assembly, side chain protection groups are removed by treatment of membranes with a deprotection cocktail (20 mL 95% trifluoroacetic acid, 3% triisopropylsilane, 2% H_2_O) for 4 hours at RT and washing (4 x DCM, 4 X EtOH, 2x H_2_O, 1 x EtOH) prior to being air dried. Peptides on arrays were derived from β-galactosidase wild-type sequence, starting from the initiator methionine. Each peptide was 13 amino acids long and neighbouring peptides were shifted along the sequence by 3 residues.

For DnaK, the peptide array was first activated in 100% MeOH for 20 s, washed for 30 min with TBS-Tween (TBS-T) and blocked for 1 hour in TBS-T with 5% non-fat dry milk and 1 hour in binding buffer (50 mM HEPES-NaOH, pH 7.5, 150 mM KCl, 10 mM MgCl_2_, 0.05% Tween, 5% milk). DnaK (1.2 µM) was incubated with the array for 1 hour at room temperature in binding buffer. Following washing in binding buffer with additional 1 mM ADP, membranes were incubated with primary antibody (1 hour at RT), washed, incubated with HRP-conjugated secondary antibody (1:10,000 dilution, 1 hour at RT) and washed again, all in binding buffer with 1 mM ADP. Finally, membranes were developed by enhanced chemiluminescence using SuperSignal West Pico PLUS Chemiluminescent Substrate (ThermoScientific). A similar protocol was followed for DnaJ with a few modifications. Binding buffer was changed to 31 mM Tris, pH 7.5, 100 mM KCl, 5 mM MgCl_2_, 0.05% Tween and 5% milk. The membrane was incubated with 0.5 µM DnaJ for 2 hours at RT and no ADP was used in buffers. As negative controls for unspecific antibody binding, the same procedure was followed on replicate membranes omitting the addition of DnaK or DnaJ.

#### Immunoblotting

Following SDS-PAGE, proteins were transferred onto a PVDF membrane using a Trans-Blot Turbo Transfer System (BioRad). Membranes were blocked in PBS-Tween with 5% non-fat milk for 1 hour at RT and incubated with appropriate primary antibodies (1:1,000 dilution in PBS-Tween with 5% non-fat milk) for 1 hour at RT. Following three washes (5 min, RT) with PBS-Tween, membranes were incubated with appropriate HRP-conjugated secondary antibodies (1:10,000 dilution in PBS-Tween with 5% non-fat milk) for 1 hour at RT and washed. Membranes were developed by enhanced chemiluminescence using SuperSignal West Pico PLUS Chemiluminescent Substrate (ThermoScientific). Antibodies used for each immunoblot are specified in figure legends and listed in the Key resource table.

### Mass spectrometry

#### Proteomic analysis of RNC composition

RNCs for proteomic analysis were purified as described above (samples labelled 1 M KOAc or high-salt) or in a modified protocol containing only one sucrose cushion centrifugation step in RNC low-salt sucrose cushion (labelled 0.1 M KOAc or low-salt). Subsequently, for each sample, 10 µg of protein estimated from the ribosome concentration (based on absorbance at 260 nm) was separated in 8 mm on NuPAGE Bis-Tris gels (1.5 mm, 12 wells, 4-12% for dataset A – Table S1; 1.0 mm, 12 wells, 12% for dataset B – Table S2 and dataset C – Table S3) followed by Quick Coomassie Stain (Neo Biotech) staining, band excision and destaining in extraction buffer (50% acetonitrile, 100 mM ammonium bicarbonate, 5 mM DTT, 16 h, 4 °C). Samples were then alkylated (40 mM chloroacetamide, 160 mM ammonium bicarbonate, 10 mM TCEP, 20 min, 70 °C), dehydrated in 100% acetonitrile, and air-dried at 37 °C, followed by digestion with trypsin (Promega). Digested peptides were processed and used to determine the protein content of each corresponding sample as described in Wales et al, 2022. In short, tryptic peptides were loaded onto Evotips (Evosep) and eluted using the 30SPD gradient via an Evosep One ^108^ HPLC with a 15 cm C18 column into a Lumos Tribrid Orbitrap mass spectrometer (ThermoScientific) via a nanospray emitter (2,200 V). Acquisition parameters were set to data-dependent mode with precursor ion spectra acquired at 120,000 resolution followed by higher energy collision dissociation. Raw files were processed in MaxQuant ^99^ and Perseus ^101^ with Uniprot *E. coli* reference proteome database and a database for common contaminants. Protein and peptide false detection rates using a decoy reverse database were set to 1%. Quantification of proteins was achieved using iBAQ (intensity-based absolute quantification) and values were plotted normalised to the average intensity of 70S ribosomal proteins. The same protocol (using 1.0 mm, 12 wells, 12% NuPAGE Bis-Tris gels) was followed when quantifying the protein content in resuspended pellets following co-sedimentation assays described above (Table S7).

#### Preparation of immobilised pepsin

Immobilisation of pepsin onto POROS 20 AL resin was carried out as previously described ^109^ with some changes. In short, porcine pepsin (Merck) was dissolved in 20 mL of 50 mM sodium citrate, pH 5 at a final concentration of 40 mg/mL. Sodium cyanoborohydrate was dissolved under an argon stream in 10 mL of 2 M Na_2_SO_4_ in 50 mM sodium citrate, pH 5 at a final concentration of 20 mg/mL and subsequently syringed into the dissolved pepsin followed by gentle mixing (10 min, RT). POROS 20 AL resin (6 g, ThermoScientific) was then added to the solution followed by mixing (10 min, RT) and dropwise addition of Na_2_SO_4_ (22 mL, 2 M), all under a constant stream of argon. After 12-16 h the reaction was quenched with dropwise addition of 1M ethanolamine in 50 mM sodium citrate, pH 5 (10 mL) followed by gentle mixing (2 h, RT). Using vacuum filtration, the resin was washed successively with 50 mM sodium citrate pH 5, 1 M NaCl in 50 mM sodium citrate, pH 5, and 50 mM sodium citrate, pH 5. The product was further washed three times with 0.08% trifluoroacetic acid (TFA), pH 2 in H_2_O by centrifugation at 7,000 g. The resin was stored at 4 °C as a 50% v/v slurry in 0.08% TFA.

#### Equilibrium HDX-MS analysis of RNCs

RNCs or RNC:chaperone complexes (prepared via a co-sedimentation assay described above) were prepared in RNC low-salt buffer at a stock concentration of 5-6 µM. Additionally, full-length native β-galactosidase, isolated chaperones and empty 70S ribosomes (NEB) were prepared as controls. Deuterium labelling was initiated by mixing 3 µL RNCs with 27 µL deuteration buffer (10 mM HEPES-NaOD, pD 7.5, 30 mM KOAc, 12 mM Mg(OAc)_2_, 1 mM DTT, RiboLock RNase inhibitor, 97% D_2_O) at 25 °C. After labelling at 25 °C for 10 or 100 seconds, the reaction was quenched with an equal volume (30 µL) of ice-cold quench buffer (100 mM sodium phosphate, pH 1.4, 4 M guanidium hydrochloride, 10 mM TCEP). The pH after quenching was 2.5. Digestion was initiated by addition of 20 µL pepsin-agarose 50% v/v bead slurry (prepared in-house as described above) equilibrated in 0.1% formic acid. Following 100 s of digestion at 10 °C with rapid mixing every 30 seconds, the sample was centrifuged (13,000 g, 15 s, 0 °C) through 0.22 µm PVDF filters (Millipore). The flow through was immediately snap frozen in liquid nitrogen for short-term storage. The same protocol was followed to prepare undeuterated controls, except the deuteration buffer was replaced by a H-based buffer (10 mM HEPES-NaOH in H_2_O, pH 7.5, 30 mM KOAc, 12 mM Mg(OAc)_2_, 1 mM DTT, RiboLock RNase inhibitor).

Frozen samples were rapidly thawed and immediately injected into an Acquity UPLC M-class system with the cooling chamber containing the chromatographic columns kept at 0 ± 0.2 °C throughout data collection. Peptic peptides were trapped and desalted for 4 minutes (200 µL/min) on a 2.1 mm X 5 mm, C4 trap column (Acquity BEH C4 Van-guard pre-column, 1.7 μm, Waters) then separated on a reverse phase Acquity UPLC HSS T3 column (1.8 µm, 1 mm X 50 mm, Waters) at a flow rate of 90 µL/min. Peptides were eluted over 25 minutes using a 3-30% gradient of acetonitrile in 0.1% formic acid. Analysis was performed using a Waters Synapt G2Si HDMS^E^ instrument in ion mobility mode, acquiring in positive ion mode over a range of 50 to 2,000 m/z with the conventional electrospray ionisation source. Calibration of the mass spectrometer was achieved using [Glu^1^]-Fibrinopeptide B (50 fmol/μL, Merck) and the instrument was operated at a source temperature of 80 °C with the capillary set to 3 kV.

MS^E^ data were processed using Protein Lynx Global Server (PLGS, Waters) to identify peptides in the undeuterated control samples using information from a non-specific cleavage of a database containing sequences of *E. coli* β-galactosidase, Trigger Factor, DnaK, DnaJ, 70S ribosomal proteins as well as porcine pepsin. PLGS search was performed using energy thresholds of low = 135 counts and elevated = 30 counts. Peptides identified by PLGS were subsequently filtered and processed in DynamX (Waters) with filters of minimum products per amino acid of 0.05 and minimum consecutive products of 1. All spectra were manually inspected, and poor-quality assignments were removed. Any peptides assigned to chaperones or β-galactosidase that were also detected in empty ribosome control samples were removed. Additionally, peptides assigned to β-galactosidase or chaperones in RNC samples, but not assigned in samples containing isolated β-galactosidase or corresponding chaperones, were also excluded. Relative deuterium uptake in Da was calculated by subtracting the centroid mass of undeuterated peptides from those of deuterated peptides. Fractional uptake was calculated by dividing the relative uptake by the theoretical maximum for each peptide, equal to n-1, where n is the peptide length excluding prolines. All HDX-MS experiments were performed in technical triplicates for each sample at each deuteration time and used to calculate the reported mean. Mean values of deuterium uptake (Table S5) are reported as relative as they are not corrected for back-exchange. Fitting of the isotopic distributions with a single or bimodal gaussian curves was done using HXExpressV3 following the published instructions ^104^.

#### Equilibrium HDX-MS of ***β***-gal truncations

Stock samples of β-galactosidase variants off ribosome were prepared at concentrations between 7-10 µM. Labelling by deuterium was initiated by a 12.5-fold dilution of 4 µL of the stock to final 50 µL using deuteration buffer (10 mM HEPES-NaOH, pD 7.5, 30 mM KOAc, 12 mM Mg(OAc)_2_, 1 mM DTT, 97% D_2_O). After labelling at 25 °C for 100 seconds, the reaction was quenched with 20 µL of 25 °C quench buffer (1 M orthophosphoric acid with pH adjusted to 2.1 using NaOH) followed by snap-freezing in liquid nitrogen. The pH after quenching was 2.35. The same protocol was followed to prepare undeuterated controls with the deuteration buffer replaced by a H-based buffer (10 mM HEPES-NaOH, pH 7.5, 30 mM KOAc, 12 mM Mg(OAc)_2_, 1 mM DTT, H_2_O).

Thawed samples were injected into an Acquity UPLC M-class system and digested on a BEH immobilized pepsin column (2.1 x 30 mm, 5 μm, Waters) held in a 20 °C chamber upstream of the cooling chamber (kept at 0 ± 0.2 °C) housing all other chromatographic columns. Pepsin digestion and trapping on a Acquity BEH C18 Van-guard trap column (Waters) was conducted for 3 min at 100 µL/min flow rate. Peptides were separated on an Acquity UPLC BEH C18 columns (1.7 μm, 100 mm x 1 mm, Waters) using a 3-35% acetonitrile gradient (16 min, 40 µL/min flow rate). The analysis was performed as described above, with two modifications. The database used for peptide searching by PLGS did not contain chaperones and 70S ribosome sequences. Additionally, more stringent filtering was applied in DynamX (minimum intensity 1,000, maximum MH+ error of 20 ppm, and minimum products per amino acid of 0.2). Values of mean deuterium uptake are reported in Table S6.

#### Crosslinking mass spectrometry

Samples for crosslinking reaction were prepared in RNC low-salt buffer at a final concentration of 1-2 µM in 50 µL volume. To detect crosslinks in chaperone-free or TF-bound RNCs (Table S4), RNCs purified from ΔTF KO cells or co-purified with TF from wild-type cells were directly used as an input into the crosslinking reaction. To detect crosslinks in DnaK-bound RNCs, RNCs purified from ΔTF cells were first incubated (30 min, 30 °C) with 8 µM DnaK, 4 µM DnaJ and 5 mM ATP. To detect crosslinks in DnaJ-bound RNCs, RNCs purified from ΔTF cells were first incubated (30 min, 30 °C) with 10 µM DnaJ. For samples containing TF-, DnaJ- and DnaK-NC crosslinks (Table S8), RNCs purified from ΔTF cells were first incubated (30 min, 30 °C) with 2 µM TF, 10 µM DnaK, 4 µM DnaJ, 5 mM ATP. The crosslinking reaction was initiated by addition of DSBU (1 mM final concentration, ThermoScientific) into the RNC-chaperone mixtures and allowed to proceed for 1 hour at 25 °C followed by quenching with 20 mM Tris-HCl, pH 7.5. Samples with excess chaperones were subsequently pelleted by centrifugation through a low-salt sucrose cushion (264,000 g, 2 h, 4 °C) and the pellet was resuspended in RNC low-salt buffer.

The crosslinked proteins were reduced with 10 mM dithiothreitol and alkylated with 50 mM iodoacetamide. They were then digested with trypsin at an enzyme-to-substrate ratio of 1:100, for 1 h at room temperature and further digested overnight at 37 °C following addition of trypsin at a ratio of 1:20. The peptide digests were then fractionated batch-wise by high pH reverse phase chromatography on micro spin TARGA C18 columns (Nest Group Inc, USA), into five fractions (10 mM NH_4_HCO_3_/10% (v/v) acetonitrile pH 8.0, 10 mM NH_4_HCO_3_/20% (v/v) acetonitrile pH 8.0, 10 mM NH_4_HCO_3_/30% (v/v) acetonitrile pH 8.0, 10 mM NH_4_HCO_3_/40% (v/v) acetonitrile pH 8.0 and 10 mM NH_4_HCO_3_/80% (v/v) acetonitrile pH 8.0). The fractions (120 µL) were evaporated to dryness in a CentriVap concentrator (Labconco, USA) prior to analysis by LC-MS/MS.

Lyophilized peptides were resuspended in 1% (v/v) formic acid and 2% (v/v) acetonitrile and analysed by nano-scale capillary LC-MS/MS using a Vanquish Neo UPLC (ThermoScientific Dionex, USA) to deliver a flow of approximately 250 nL/min. A PepMap Neo C18 5 μm, 300 μm x 5 mm nanoViper (ThermoScientific Dionex, USA) trapped the peptides before separation on a 50 cm EASY-Spray column (50 cm x 75 µm ID, PepMap C18, 2 µm particles, 100 Å pore size: ThermoScientific, USA). Peptides were eluted with a gradient of acetonitrile over 90 minutes. The analytical column outlet was directly interfaced via a nano-flow electrospray ionisation source, with a quadrupole Orbitrap mass spectrometer (Orbitrap Exploris 480, ThermoScientific, USA). MS data were acquired in data-dependent mode using a top 10 method, where ions with a precursor charge state of 1+ and 2+ were excluded. High-resolution full scans (R=60000, m/z 380-1800) were recorded in the Orbitrap followed by higher energy collision dissociation (HCD) (stepped collision energy 30 and 32% Normalized Collision Energy) of the 10 most intense MS peaks. The fragment ion spectra were acquired at a resolution of 30,000 and a dynamic exclusion window of 20 secs was applied.

Data analysis was performed as described previously ^110^ with small changes. Xcalibur raw files were converted into the MGF format using Proteome Discoverer version 2.3 (ThermoScientific) and used directly as input files for MeroX ^98^. Searches were performed against an ad hoc protein database containing the sequences of the proteins in the complex and a set of randomized decoy sequences generated by the software. The following parameters were set for the searches: maximum number of missed cleavages: 3; targeted residues K, S, Y and T; minimum peptide length 5 amino acids; variable modifications: carbamidomethylation of cysteine (mass shift 57.02146 Da), methionine oxidation (mass shift 15.99491 Da); DSBU modified fragments: 85.05276 Da and 111.03203 Da (precision: 5 ppm MS and 10 ppm MS/MS); False Discovery Rate cut-off: 5%. Finally, each fragmentation spectrum was manually inspected and validated. Crosslinks to non-native N-terminal sequences (present in purified proteins because of purification tag cleavage) or crosslinks with score below 50 were not considered, unless explicitly highlighted. PyXlinkViewer ^103^ plugin and xiVIEW online tool ^100^ were used to visualise crosslinks on 3D structures and linear domain diagrams, respectively.

## QUANTIFICATION AND STATISTICAL ANALYSIS

Band intensities on SDS-PAGE gels were quantified in Fiji. All statistical analyses were performed in GraphPad Prism 9. Quantification of iBAQ values was performed in MaxQuant. Gaussian fits of HDX-MS spectra were performed in HXExpressV3^104,111^. Hydrophobicity values of solved protein structures were calculated using ChimeraX Molecular Lipophilicity Potential ^91^.

## ACKNOWLEDGEMENTS

We thank F.U. Hartl (MPI Biochemistry) for the plasmids encoding TF, DnaK, DnaJ and GrpE; John Christodoulou (University College London) for the ΔTF *E. coli* strain; Stephane Mouilleron (Francis Crick institute) for 3C protease and TEV protease; Simone Kunzelmann and Laura Masino (Francis Crick Institute Structural Biology STP) for advice on the chaperone binding experiments; Christelle Soudy (Francis Crick Institute Chemical Biology STP) for help with preparing immobilised pepsin; Thomas Wales and John Engen (Northeastern University) for advice on the HDX-MS experiments; and all members of the Protein Biogenesis Lab (Francis Crick Institute) for useful discussion. DB’s work is supported by the Francis Crick Institute which receives its core funding from Cancer Research UK (FC001985), the UK Medical Research Council (FC001985), and the Wellcome Trust (FC001985). For the purpose of Open Access, the author has applied a CC BY public copyright licence to any Author Accepted Manuscript version arising from this submission.

## AUTHOR CONTRIBUTIONS

A.R. performed most of the experiments, analysed the data and wrote the manuscript together with D.B. S.L.M. and J.M.S. collected and processed the XL-MS data and together with D.B. provided technical assistance and training for HDX-MS experiments. S.S. purified TF_ΔPPD_ and full-length β-galactosidase proteins. G.A.P. and S.S contributed to optimising the HDX-MS workflows and prepared pepsin-coupled beads. S.H. performed the proteomic analysis of RNC composition. D.J. and S.R. prepared the peptide arrays. S.K. contributed to designing the RNC isolation strategy and prepared the GFP-clamp resin. D.B. conceived and supervised the project.

## DECLARATION OF INTERESTS

The authors do not declare any interests.

## SUPPLEMENTAL INFORMATION TITLES AND LEGENDS

**Figure S1. Characterisation of stalled RNCs, related to Figure 1**

**(A)** Mean intensity-based absolute quantification (iBAQ) values of ribosomal 30S (top) and 50S (bottom) proteins, normalised to the average iBAQ of all ribosomal proteins in each corresponding RNC sample purified via two high-salt sucrose cushions. Values for each ribosomal protein are plotted as the mean of 3 technical replicates for each of 14 different RNC complexes with the associated SD.

**(B)** iBAQ values for β-gal in each RNC sample purified via one low-salt sucrose cushion. Values are normalised to the average iBAQ of all ribosomal proteins in each sample, and corrected for the length of the NC in each RNC. Value of 100% would correspond to a1:1 NC:ribosome ratio. Error-bars represent SD of 1-4 technical replicates.

**(C)** 70S-normalised iBAQ values for the most abundant proteins copurifying with RNCs after two high-salt sucrose cushions. Ribosomal proteins and β-gal were not included in this plot. Error-bars correspond to the SD of 3 technical replicates.

**(D)** Coomassie-stained SDS-PAGE gel showing the ratio of tRNA-linked (+tRNA) and tRNA-free (-tRNA) NC_1-646_ upon various treatments of RNC_1-646_. Treatments included incubation at 37 °C for 1 h, five subsequent freeze-thaw cycles (5xFT), and 37 °C incubation with additional 1.5 M hydroxylamine (HA), 1 mM puromycin dihydrochloride (Puro) or 50 mM EDTA with 50 μg/mL RNaseA (RNase). Ribosomal protein S1 is shown as a control.

**(E)** β-gal enzyme activity of select RNCs, empty 70S ribosomes and isolated native β-gal. Error-bars represent the SD of 3 technical replicates.

**(F)** Coomassie-stained SDS-PAGE of the ribosomal fraction after a co-sedimentation assay of a mixture of 70S ribosomes with TF, centrifuged through either a 0.1 M or 1 M KOAc sucrose cushion. The band corresponding to TF is indicated.

**(G)** Coomassie-stained SDS-PAGE gel of the ribosomal fraction after a co-sedimentation assay of a mixture of 70S ribosomes or RNC_1-646_, with either wild-type TF (TF_WT_) or TF with mutated ribosome-binding site (TF_ΔRBS_) centrifuged through a 1 M KOAc sucrose cushion. The band corresponding to TF is indicated.

**(H)** Fluorescence intensity of 2 μM BADAN-labelled TF_R14C_ (TF-BADAN) measured during the addition of 2 μM RNC_1-646_. 10 µM unlabelled TF_R14C_ is added where indicated. BADAN fluorescence is quenched upon TF binding to RNCs, and relieved by competition with unlabelled TF.

**(I)** As in (H), performed using 2 μM BADAN-labelled TF_R14C_ with a mutated ribosome-binding site (ΔRBS-BADAN).

**(J)** Equilibrium titration of RNC_1-646_ against 5 nM BADAN-labelled TF_R14C_. Samples were incubated for 15 min at 25 °C before recording BADAN fluorescence. Values correspond to the mean of 3 independent measurements of each sample. Fitting a one-site specific binding model (GraphPad Prism) yielded a K_D_ of 21 nM with a 95% CI of 14 to 32 nM.

**(K)** Coomassie-stained SDS-PAGE showing the amount of TF that co-purifies with 10 µg of indicated RNCs using one low-salt (0.1 M KOAc) or two high-salt (1 M KOAc) sucrose cushion steps in the purification procedure. These gels are cropped from those shown in Fig 1D and FigS5A.

**Figure S2. Determinants of TF binding to RNCs, related to Figure 2**

**(A)** Domain organisation and corresponding 2D schematic of wild-type Trigger factor (TF_WT_), Trigger factor with a mutated RBS (TF_ΔRBS_), Trigger factor with a deleted PPD (TF_ΔPPD_), isolated His_6_-SUMO-tagged TF RBD (RBD_N-tag_) and the Trigger factor variant with two RBDs (TF_2xRBD_). In TF_2xRBD_, only the N-terminal RBD encodes the wild-type RBS. Sequences of all visualised proteins are listed in Table S11.

**(B)** Coomassie-stained SDS-PAGE showing the bands corresponding to TF and ribosomal protein S1 in the resuspended ribosomal pellet from co-sedimentation assays of empty ribosomes (70S) or RNCs purified from ΔTF cells, incubated with either WT TF (TF_WT_) or TF without the PPD (TF_ΔPPD_). The KOAc salt content in the sucrose cushion was either 0.1 M or 1 M as indicated.

**(C)** Top: Coomassie-stained SDS-PAGE of the resuspended ribosomal pellet from co-sedimentation assays of empty ribosomes (70S) or RNCs, incubated with RBD_N-tag_. The KOAc salt content in the sucrose cushion was either 0.1 M or 1 M as indicated. Bands corresponding to NCs (*) and the RBD_N-tag_ (●) are indicated. Bottom: Immunoblot of a replicate SDS-PAGE gel probed against TF, showing the presence of RBDN-tag.

**(D)** Experimentally determined structure of TF monomer (left, PDB: 1W26) and pre dicted structure of TF_2xRBD_ (right, AlphaFold2.0) with the ribosome-binding domain(s) coloured grey and the PPD and CTD in green.

**(E)** Plots of the difference in deuterium uptake after 10 s (green) or 100 s (black) deuteration between isolated TF and TF bound to 70S ribosomes, RNC_1-646_, RNC_1-687_ and RNC_1-1014_. Values are plotted for individual peptides covering TF detected in the HDX-MS dataset. Negative values indicate less deuteration of a peptide in 70S/RNC-bound TF relative to isolated TF. Uptake values are listed in Table S5.

**Figure S3. TF interaction with nascent chains, related to Figure 3**

**(A)** Plot showing the number of residues between the C-terminus of each RNC and the most C-terminal NC residue detected to crosslink to TF CTD (left) or PPD (right). The vertical line and grey shading indicates the number of residues required to span the ribosome exit tunnel (assumed to be 30 aa).

**(B)** Plot showing the maximum distance (in aa) between NC residues found to crosslink the TF CTD (left) or PPD (right).

**(C)** Crosslinks between TF and RNCs visualised on TF and β-gal domain diagrams. Crosslinks are colour-coded based on the crosslinked TF domain (RBD – red, PPD – green, CTD – blue).

**(D)** Histograms showing the frequency distribution of NC residues based on the fractional deuterium uptake difference between RNCs and native β-gal for RNC_1-646_ (left) and RNC_1-687_ (right), in isolation (black) or bound by TF (grey) at deuteration time 10 s. Fractional uptake values are listed in Table S5.

**(E)** Position of β-gal residues forming intramolecular crosslinks upon DSBU crosslinking of β-gal RNCs in complex with TF, as listed in Table S4. Residues close in amino acid sequence were grouped and coloured accordingly. Numbers above the domain diagram indicate the RNCs for which the crosslinks were detected. E.g. RNC_305_ and RNC_352_ form intra-NC crosslinks connecting the N-terminal region and D1.

**(F)** Amount of undigested NCs present at different time points after limited proteinase K digestion of indicated RNC in isolation (-TF, black) or upon incubation with Trigger factor (TF_WT_, green). Error-bars correspond to SD of values from 3 independent reactions.

**(G)** Difference in deuterium uptake (after 10 s deuteration) between RNC_1-646_ with or without bound TF. Values are plotted for individual peptides covering NC_1-646_ with region 504-535 highlighted. Positive values indicate deprotection of the NC induced by TF.

**(H)** Mass spectral envelopes corresponding to peptide 504-535 of β-gal NC_1-687_, detected in HDX-MS data of isolated or TF bound (+TF) RNC_1-687_ samples. The isotopic distributions were fit to single or bimodal gaussian curves (red and green) using HXExpressV3 ^104^.

**Figure S4. Effects of destabilising mutations and truncations on β-gal, related to Figure 4**

**(A)** Coomassie-stained SDS-PAGE of the resuspended ribosomal pellet from co-sedimentation assays of RNCs (1-510, 1-510_+11G/S_ and 1-510_+50G/S_) purified from WT *E. coli* and incubated with additional TF *in vitro*. The KOAc salt content in the sucrose cushion was 0.1 or 1 M as indicated. The expected position of TF is indicated, and bands corresponding to NCs are labelled (*).

**(B)** Top: Structure of β-gal (PDB: 6CVM) with residues I141, V567 and L652 shown as red spheres. Bottom: Coomassie-stained SDS-PAGE of the soluble (S) and pellet/insoluble (P) fraction of a cell lysate from *E. coli* cells overexpressing truncated β-gal constructs 1-490_WT_, 1-490_I141N_, 1-725_V567D_ and 1-725_L652D_.

**(C)** Coomassie-stained SDS-PAGE gels of the soluble (top, supernatant) and insoluble (bottom, pellet) fractions of a cell lysate from *E. coli* cells overexpressing truncated β-gal constructs. Full-length (FL) β-gal is included as a control in the last lane.

**(D)** Left: histograms showing the frequency distribution of residues based on the fractional deuterium uptake difference between truncated chains (1-332, 1-440, 1-490, 1-725 and 1-725_V567D_) and full-length β-gal. Right: Plot of the difference in deuterium uptake (after 100 s deuteration) between truncated chains and full-length β-gal. Values are plotted for individual peptides covering β-gal in each chain, and regions that are destabilised in multiple chains are highlighted in red. Higher values indicate more deuteration of peptides in truncated chains relative to full-length β-gal. Uptake values are listed in Table S6.

**Figure S5. Determinants of DnaK/J binding to RNCs, related to Figure 5**

**(A)** Top: Coomassie-stained SDS-PAGE of β-gal RNCs purified via one 0.1 M KOAc sucrose cushion from WT BL21(DE3) *E. coli*. Bands corresponding to nascent chains (*) migrate slower than expected based on protein molecular weight, due to the covalently bound tRNA (∼20 kDa). Some RNCs co-purify with Trigger factor (TF, ∼50 kDa) at close to 1:1 stoichiometry. The band corresponding to 3C protease (∼40 kDa) used in purification (Fig 1C) is indicated. All other major bands present in the purified RNC samples correspond to 70S ribosomal proteins including S1 (∼70 kDa) Bottom: immunoblot of a replicate SDS-PAGE gel probed against DnaK, DnaJ, GroEL and ribosomal protein S2.

**(B)** Intensity-based absolute quantification (iBAQ) values for DnaK (black) and DnaJ (grey) normalised to the average iBAQ of all ribosomal proteins in each corresponding RNC sample, purified via one low-salt sucrose cushion.

**(C)** Mean iBAQ for DnaJ present in the resuspended ribosomal pellet from high-salt co-sedimentation assays of RNC_1-646_, incubated with either DnaJ alone (+J) or with DnaJ and excess DnaK (+KJ), both in the presence of ATP. The iBAQ values were normalised to the average iBAQ value of all 70S ribosomal proteins. Error-bars correspond to the SD from 3 independent co-sedimentation assays.

**(D)** Coomassie-stained SDS-PAGE of the resuspended ribosomal pellet from high-salt co-sedimentation assays of 70S ribosomes incubated with either DnaK (+K), DnaJ (+J), or both DnaK and DnaJ (+KJ) in the presence and absence of ATP as indicated, as well as with additional GrpE (+KJ+E). The bands corresponding to ribosomal protein S1 are indicated.

**(E)** Coomassie-stained SDS-PAGE gel of the resuspended ribosomal pellet from high-salt co-sedimentation assays of RNC_1-646_ incubated with either DnaK and DnaJ (+KJ) or with excess (25 µM) GrpE (+KJ+↑E), both in the presence of ATP. The bands corresponding to DnaK and DnaJ are indicated.

**(F)** Coomassie-stained SDS-PAGE of the resuspended ribosomal pellet from high-salt co-sedimentation assays of RNCs incubated, in the presence of ATP, with either TF (+TF), DnaK/DnaJ (+KJ), or all three factors added in the specified order (+KJ+TF or +TF+KJ). The bands corresponding to DnaK and TF are indicated. The first gel shows the migration of purified TF, GrpE, DnaJ and DnaK during SDS-PAGE, for reference.

**(G)** Coomassie-stained SDS-PAGE of the resuspended ribosomal pellet from high-salt co-sedimentation assays of RNCs incubated with DnaK and DnaJ in the presence of ATP. The bands corresponding to DnaK are indicated.

**(H)** Coomassie-stained SDS-PAGE of the resuspended ribosomal pellet from high-salt co-sedimentation assays of RNCs incubated with either TF (+TF) or DnaK with DnaJ (+KJ) or all three factors (+KJ+TF), all in the presence of ATP. The bands corresponding to DnaK and TF are indicated.

**(I)** Peptide array developed with ECL after incubation with DnaK and appropriate antibodies. Each position on the grid contains a 13-residue peptide derived from the wild-type β-gal sequence, starting from the initiator methionine (numbered as residue 0) at position A1. The sequence of peptides is shifted by 3 residues between neighbouring spots. The peptide in position R21 is the NRLLLTG peptide, serving as a positive control. Sites identified as potential DnaK binding sites are boxed in red and labelled with a letter (A, C, E, F, J, K, P) followed by the exact sequence.

**(J)** Structure of truncated β-galactosidase (PDB: 6CVM) coloured as in Fig 5H, with assembly interfaces indicated using orange lines.

**(K)** Coomassie-stained SDS-PAGE of the resuspended ribosomal pellet from high-salt co-sedimentation assays of 70S ribosome and RNC_1-646_ variants (RNC with a loop deletion - Δ504-532, RNC with limbo-predicted DnaK binding site H deletion – ΔH and wild-type RNC – WT) incubated with either TF (+TF), DnaJ (+J) or DnaK with DnaJ (+KJ), all in the presence of ATP. The bands corresponding to DnaK, DnaJ and TF are indicated.

**Figure S6. DnaK/J interaction with nascent chains, related to Figure 5 and Figure 6**

**(A)** Difference in deuterium uptake after 10 s (purple) or 100 s (black) deuteration, between isolated DnaK and DnaK bound to RNC_1-646_ (top) or RNC_1-1014_ (bottom). Values are plotted for individual peptides covering DnaK detected in the HDX-MS dataset. Negative values indicate less deuteration of a peptide in RNC-bound DnaK relative to isolated DnaK. Uptake values are listed in Table S5.

**(B)** Number of unique crosslinks detected between DnaJ and each group of residues (as in Fig 3A) on the specified NC.

**(C)** Number of DSBU-reactive residues (K, S, T or Y) in each domain of DnaJ.

**(D)** Difference in deuterium uptake after 10 s (blue) or 100 s (black) deuteration, between isolated DnaJ and DnaJ bound to RNC_1-646_. Values are plotted for individual peptides covering DnaJ detected in the HDX-MS dataset. Negative values indicate less deuteration of a peptide in RNC-bound DnaJ relative to isolated DnaJ.

**(E)** Left: DnaJ monomer structure (AF-P08622-F1), indicating site A (120-140, yellow) and site B (214-244, blue). Right: AlphaFold 2.0 multimer-predicted structure DnaJ monomer in complex with peptide GWLYEIS. The top 4 most confidently predicted structures are overlayed.

**(F)** Structure of CTD I (left) and CTD II (right) of DnaJ (AF-P08622-F1), with regions protected from deuterium uptake in RNC-bound DnaJ relative to isolated DnaJ coloured blue, and regions not covered by any detected peptides coloured grey.

**(G)** Sequence alignment of structurally homologous sites on CTD I and CTD II of DnaJ.

**(H)** Domain diagram of DnaJ showing intra-DnaJ crosslinks which were detected in DnaJ bound to RNC_1-646_. Crosslinks between equivalent residues in two different DnaJ molecules - are shown in red.

**(I)** Peptide arrays developed with ECL after incubation with DnaJ and appropriate antibodies (left) or only with antibodies (right). Each position contains a 13-residue peptide derived from the wild-type β-gal sequence, starting from the initiator methionine (numbered as residue 0) at position A1. The sequence of peptides is shifted by 3 residues between neighbouring spots. Peptides identified as potential DnaJ binding sites are circled in red.

**Figure S7. Cooperation and competition of DnaK, DnaJ and TF on RNCs, related to Figure7**

**(A)** Domain schematic of RNC_1-646_ (top) and RNC_1-687_ (bottom), showing the position of sites affected by indicated chaperones based on HDX-MS experiments described in Fig 3, 5 and 6.

**(B)** Coomassie-stained SDS-PAGE of the resuspended ribosomal pellet from high-salt co-sedimentation assays of indicated RNCs incubated with either DnaJ (J), TF, or both (J+TF). The bands corresponding to DnaJ and TF are indicated.

**(C)** Close-up of the 70S ribosome exit port (red), with protein L29 coloured green and residue K4 in orange (PDB: 7D80). DnaJ, TF and β-gal nascent chains in RNC complexes formed crosslinks with a lysine on a tryptic peptide AKELR. Note that this peptide is present in both ribosomal proteins L29 (_3_AKELR_7_) and L17 (_41_AKELR_45_). However, only K4 in L29 is solvent-accessible.

**(D)** Amount of undigested NC_1-646_ present at different time points after limited proteinase K digestion of RNC_1-646_, in isolation (-DnaJ, black) or upon incubation with DnaJ (+DnaJ, blue). Error-bars correspond to the SD of 3 independent reactions.

**(E)** Coomassie-stained SDS-PAGE of the resuspended ribosomal pellet from high-salt co-sedimentation assays of RNC_1-180_, incubated with either wild-type TF (+TF_WT_), TF without the PPD (+TF_ΔPPD_), DnaK and DnaJ (+KJ) or combination of DnaK and DnaJ with either TF variant (+KJ+TF_WT_ or +KJ+TF_ΔPPD_), all in the presence of ATP. The bands corresponding to DnaK and TF variants are indicated.

**(F)** Schematic showing the position of intra-NC (black) and chaperone-NC (grey) crosslinks detected in RNCs 1-646 (left), 1-900 (middle) and 1-1014 (right), with or without bound chaperones (TF, DnaJ or DnaK). Samples were prepared under four different conditions. From top to bottom: RNCs purified from ΔTF cells, RNCs purified from WT cells; RNCs purified from ΔTF cells and incubated with DnaJ; RNCs purified from ΔTF cells then incubated with DnaK, DnaJ and ATP. Crosslinked RNCs incubated with excess chaperones were subjected to an additional centrifugation through a high-salt sucrose cushion to remove unbound factors before trypsin digestion and MS analysis.

**Table S1.** Proteomic analysis of RNC composition in dataset A, related to Figure 1. List of detected proteins and corresponding intensities

**Table S2.** Proteomic analysis of RNC composition in dataset B, related to Figure 1. List of detected proteins and corresponding intensities

**Table S3.** Proteomic analysis of RNC composition in dataset C, related to Figure 1 and S5. List of detected proteins and corresponding intensities

**Table S4.** List of residues crosslinked in RNC samples purified from ΔTF or WT *E.coli*, related to Figure 2, 3 and 7

**Table S5.** Summary of differential HDX-MS of β-galactosidase RNCs with and without bound chaperones, related to Figure 2, 3, 5, 6 and 7

**Table S6.** Summary of differential HDX-MS of β-galactosidase truncations compared to native full-length β-galactosidase, related to Figure 4

**Table S7.** Proteomic analysis of resuspended pellets from co-sedimentation assays of RNCs with chaperones, related to Figure 5 and 7

**Table S8.** List of residues crosslinked in RNC samples purified from ΔTF *E. coli* and incubated with recombinant chaperones, related to Figure 5, 6 and 7

**Table S9.** LIMBO-predicted DnaK binding sites on β-galactosidase, related to Figure 5

**Table S10.** List of recombinant DNA used in this study, related to STAR Methods

**Table S11.** List of protein sequences of recombinant proteins used in this study, related to STAR Methods

## DATA AND CODE AVAILABILITY

The mass spectrometry proteomics data have been deposited to the ProteomeXchange Consortium via the PRIDE ^112^ partner repository with the following dataset identifiers:

RNC composition: PXD048645

HDX-MS of RNCs: PXD048642

HDX-MS of truncated β-gal: PXD048638

XL-MS: PXD048623

## Notes

### Competing Interest Statement

The authors have declared no competing interest.

