## Supplementary Figures for "Mechanism of chaperone coordination during cotranslational protein folding in bacteria"

#### Figure S1

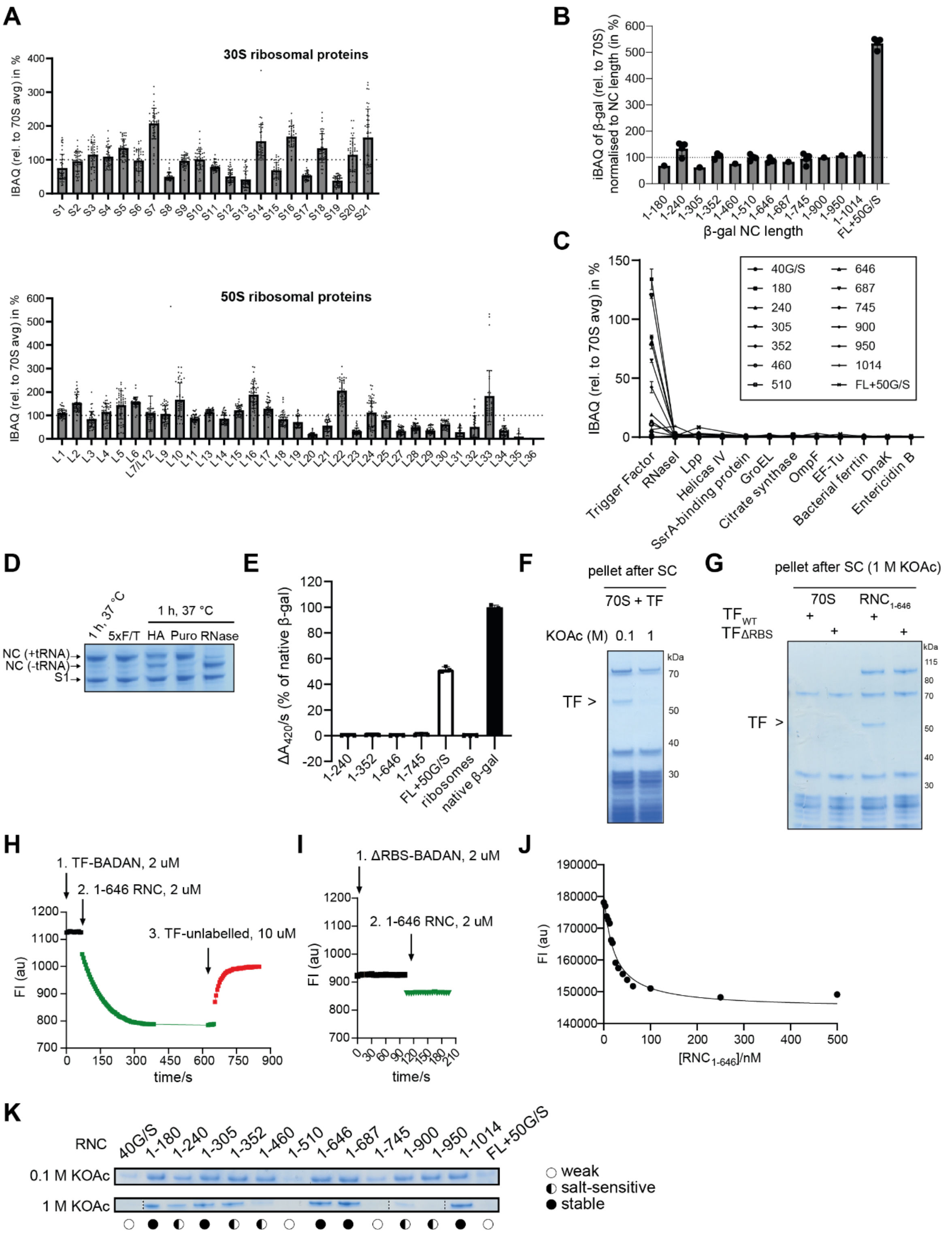

**Figure S1. Characterisation of stalled RNCs, related to Figure 1**

**(A)** Mean intensity-based absolute quantification (iBAQ) values of ribosomal 30S (top) and 50S (bottom) proteins, normalised to the average iBAQ of all ribosomal proteins in each corresponding RNC sample purified via two high-salt sucrose cushions. Values for each ribosomal protein are plotted as the mean of 3 technical replicates for each of 14 different RNC complexes with the associated SD.

**(B)** iBAQ values for β-gal in each RNC sample purified via one low-salt sucrose cushion. Values are normalised to the average iBAQ of all ribosomal proteins in each sample, and corrected for the length of the NC in each RNC. Value of 100% would correspond to a1:1 NC:ribosome ratio. Error-bars represent SD of 1-4 technical replicates.
**(C)** 70S-normalised iBAQ values for the most abundant proteins copurifying with RNCs after two high-salt sucrose cushions. Ribosomal proteins and β-gal were not included in this plot. Error-bars correspond to the SD of 3 technical replicates.
**(D)** Coomassie-stained SDS-PAGE gel showing the ratio of tRNA-linked (+tRNA) and tRNA-free (-tRNA) $NC_{1-646}$ upon various treatments of $RNC_{1-646}$. Treatments included incubation at 37 °C for 1 h, five subsequent freeze-thaw cycles (5xFT), and 37 °C incubation with additional 1.5 M hydroxylamine (HA), 1 mM puromycin dihydrochloride (Puro) or 50 mM EDTA with 50 µg/mL RNaseA (RNase). Ribosomal protein S1 is shown as a control.
**(E)** β-gal enzyme activity of select RNCs, empty 70S ribosomes and isolated native β-gal. Error-bars represent the SD of 3 technical replicates.
**(F)** Coomassie-stained SDS-PAGE of the ribosomal fraction after a co-sedimentation assay of a mixture of 70S ribosomes with TF, centrifuged through either a 0.1 M or 1 M KOAc sucrose cushion. The band corresponding to TF is indicated.
**(G)** Coomassie-stained SDS-PAGE gel of the ribosomal fraction after a co-sedimentation assay of a mixture of 70S ribosomes or $RNC_{1-646}$, with either wild-type TF ($TF_{WT}$) or TF with mutated ribosome-binding site ($TF_{\Delta RBS}$) centrifuged through a 1 M KOAc sucrose cushion. The band corresponding to TF is indicated.
**(H)** Fluorescence intensity of 2 µM BADAN-labelled $TF_{R14C}$ (TF-BADAN) measured during the addition of 2 µM $RNC_{1-646}$. 10 µM unlabelled $TF_{R14C}$ is added where indicated. BADAN fluorescence is quenched upon TF binding to RNCs, and relieved by competition with unlabelled TF.
**(I)** As in (H), performed using 2 µM BADAN-labelled $TF_{R14C}$ with a mutated ribosome-binding site (ΔRBS-BADAN).
**(J)** Equilibrium titration of $RNC_{1-646}$ against 5 nM BADAN-labelled $TF_{R14C}$. Samples were incubated for 15 min at 25 °C before recording BADAN fluorescence. Values correspond to the mean of 3 independent measurements of each sample. Fitting a one-site specific binding model (GraphPad Prism) yielded a $K_D$ of 21 nM with a 95% CI of 14 to 32 nM.
**(K)** Coomassie-stained SDS-PAGE showing the amount of TF that co-purifies with 10 µg of indicated RNCs using one low-salt (0.1 M KOAc) or two high-salt (1 M KOAc) sucrose cushion steps in the purification procedure. These gels are cropped from those shown in Fig 1D and FigS5A.

#### Figure S2

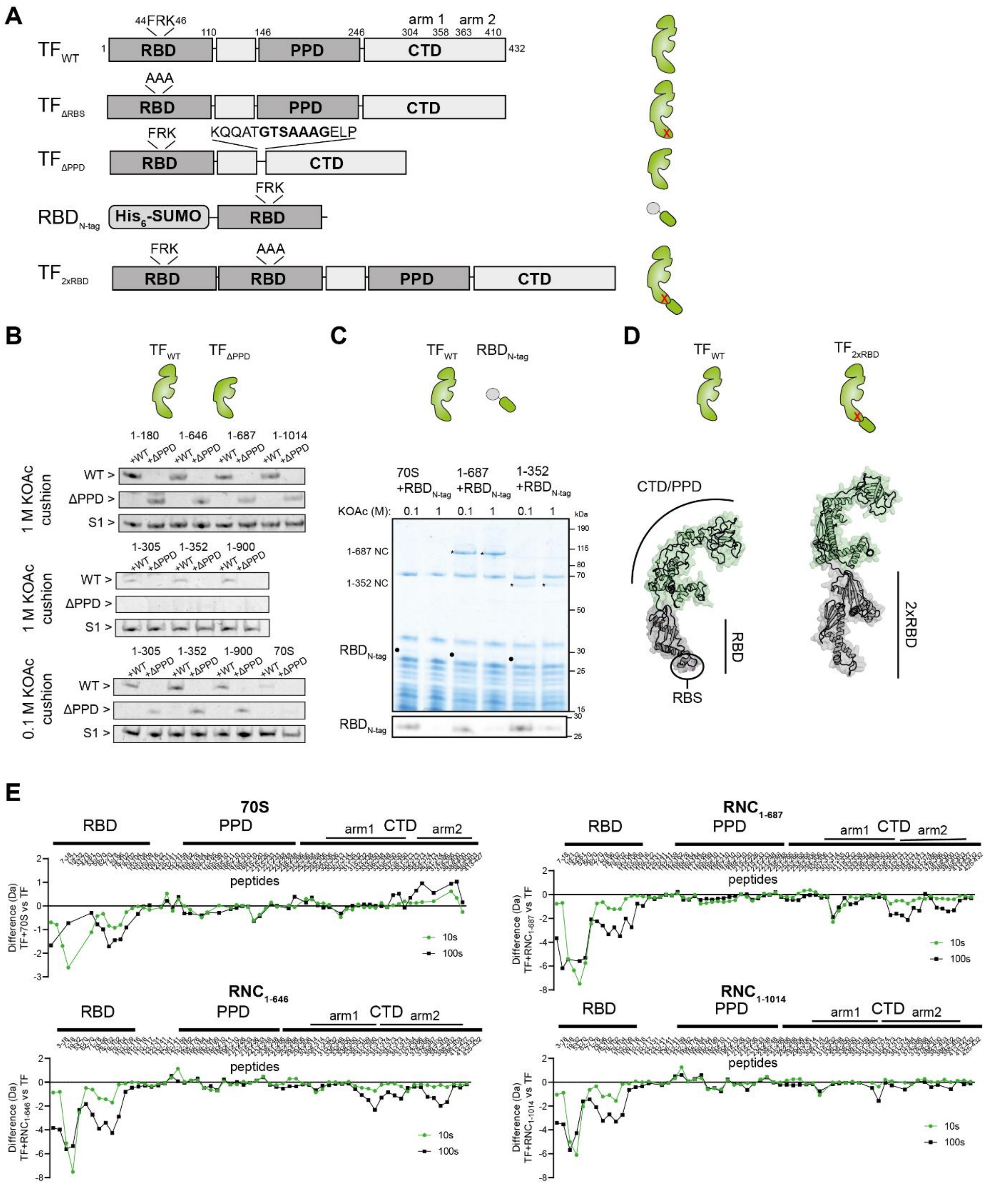


**Figure S2. Determinants of TF binding to RNCs, related to Figure 2**

**(A)** Domain organisation and corresponding 2D schematic of wild-type Trigger factor ($TF_{WT}$), Trigger factor with a mutated RBS ($TF_{\Delta RBS}$), Trigger factor with a deleted PPD ($TF_{\Delta PPD}$), isolated $His_6$-SUMO-tagged TF RBD ($RBD_{N\text{-}tag}$) and the Trigger factor variant with two RBDs ($TF_{2xRBD}$). In $TF_{2xRBD}$, only the N-terminal RBD encodes the wild-type RBS. Sequences of all visualised proteins are listed in Table S11.

**(B)** Coomassie-stained SDS-PAGE showing the bands corresponding to TF and ribosomal protein S1 in the resuspended ribosomal pellet from co-sedimentation assays of empty ribosomes (70S) or RNCs purified from ΔTF cells, incubated with either WT TF ($TF_{WT}$) or TF without the PPD ($TF_{\Delta PPD}$). The KOAc salt content in the sucrose cushion was either 0.1 M or 1 M as indicated.
**(C)** Top: Coomassie-stained SDS-PAGE of the resuspended ribosomal pellet from co-sedimentation assays of empty ribosomes (70S) or RNCs, incubated with $RBD_{N\text{-}tag}$. The KOAc salt content in the sucrose cushion was either 0.1 M or 1 M as indicated. Bands corresponding to NCs (*) and the $RBD_{N\text{-}tag}$ (●) are indicated. Bottom: Immunoblot of a replicate SDS-PAGE gel probed against TF, showing the presence of $RBD_{N\text{-}tag}$.
**(D)** Experimentally determined structure of TF monomer (left, PDB: 1W26) and predicted structure of $TF_{2xRBD}$ (right, AlphaFold2.0) with the ribosome-binding domain(s) coloured grey and the PPD and CTD in green.
**(E)** Plots of the difference in deuterium uptake after 10 s (green) or 100 s (black) deuteration between isolated TF and TF bound to 70S ribosomes, $RNC_{1\text{-}646}$, $RNC_{1\text{-}687}$ and $RNC_{1\text{-}1014}$. Values are plotted for individual peptides covering TF detected in the HDX-MS dataset. Negative values indicate less deuteration of a peptide in 70S/RNC-bound TF relative to isolated TF. Uptake values are listed in Table S5.

### Figure S3

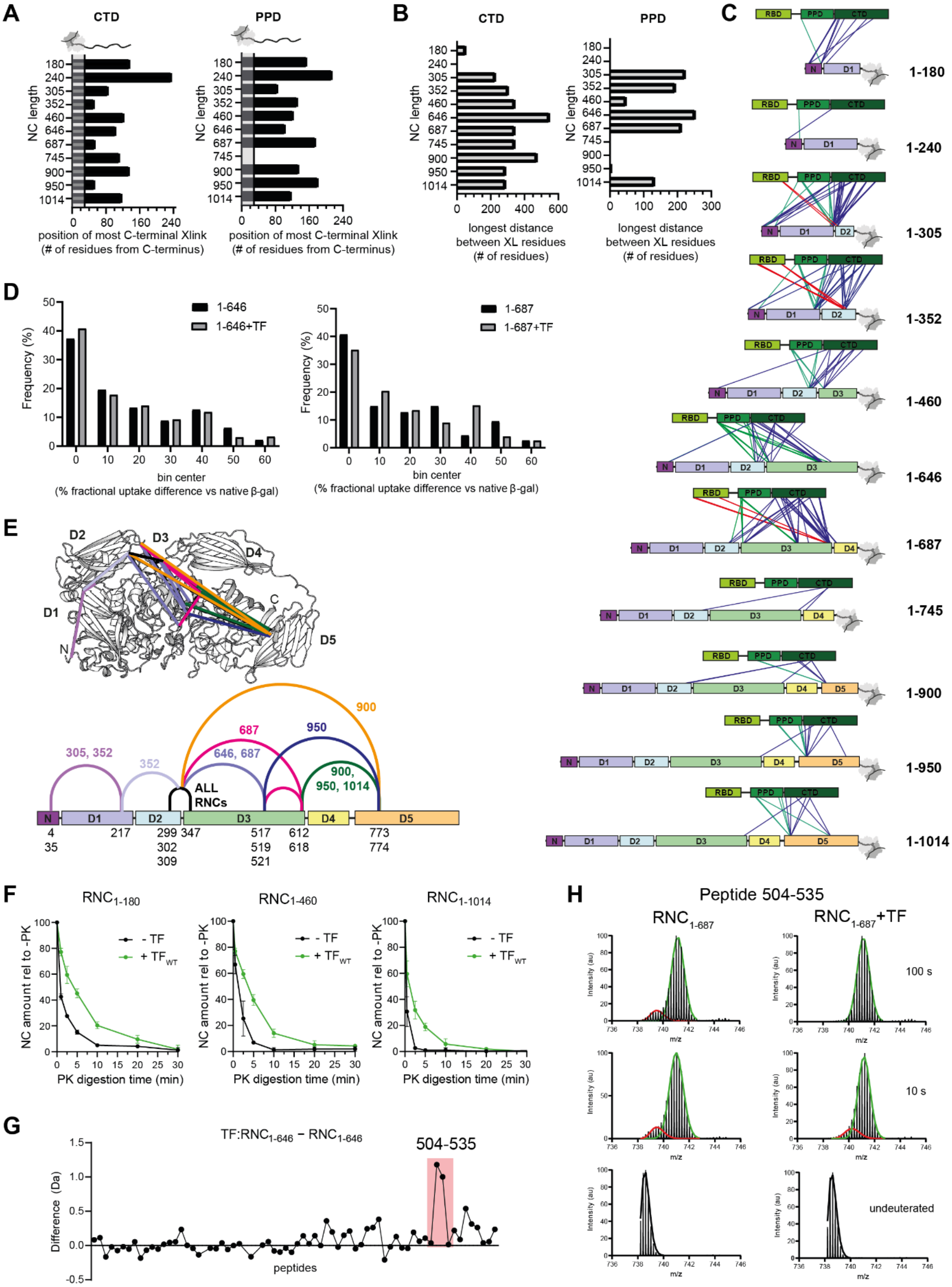
A
CTD
PPD
NC length
position of most C-terminal Xlink
(# of residues from C-terminus)
B
longest distance
between XL residues
(# of residues)
C
RBD
PPD
CTD
1-180
1-240
1-305
1-352
1-460
1-646
1-687
1-745
1-900
1-950
1-1014
D
1-646
1-646+TF
1-687
1-687+TF
Frequency (%)
bin center
(% fractional uptake difference vs native β-gal)
E
D1
D2
D3
D4
D5
ALL RNCs
F
RNC1-180
RNC1-460
RNC1-1014
- TF
+ TFWT
NC amount rel to -PK
PK digestion time (min)
G
TF:RNC1-646 – RNC1-646
504-535
Difference (Da)
peptides
H
Peptide 504-535
RNC1-687
RNC1-687+TF
100 s
10 s
undeuterated
Intensity (au)
m/z

**Figure S3. TF interaction with nascent chains, related to Figure 3**
**(A)** Plot showing the number of residues between the C-terminus of each RNC and the most C-terminal NC residue detected to crosslink to TF CTD (left) or PPD (right). The vertical line and grey shading indicates the number of residues required to span the ribosome exit tunnel (assumed to be 30 aa).
**(B)** Plot showing the maximum distance (in aa) between NC residues found to crosslink the TF CTD (left) or PPD (right).
**(C)** Crosslinks between TF and RNCs visualised on TF and β-gal domain diagrams. Crosslinks are colour-coded based on the crosslinked TF domain (RBD – red, PPD – green, CTD – blue).
**(D)** Histograms showing the frequency distribution of NC residues based on the fractional deuterium uptake difference between RNCs and native β-gal for $RNC_{1\text{-}646}$ (left) and $RNC_{1\text{-}687}$ (right), in isolation (black) or bound by TF (grey) at deuteration time 10 s. Fractional uptake values are listed in Table S5.
**(E)** Position of β-gal residues forming intramolecular crosslinks upon DSBU crosslinking of β-gal RNCs in complex with TF, as listed in Table S4. Residues close in amino acid sequence were grouped and coloured accordingly. Numbers above the domain diagram indicate the RNCs for which the crosslinks were detected. E.g. $RNC_{305}$ and $RNC_{352}$ form intra-NC crosslinks connecting the N-terminal region and D1.
**(F)** Amount of undigested NCs present at different time points after limited proteinase K digestion of indicated RNC in isolation (-TF, black) or upon incubation with Trigger factor ($TF_{WT}$, green). Error-bars correspond to SD of values from 3 independent reactions.
**(G)** Difference in deuterium uptake (after 10 s deuteration) between $RNC_{1\text{-}646}$ with or without bound TF. Values are plotted for individual peptides covering $NC_{1\text{-}646}$ with region 504-535 highlighted. Positive values indicate deprotection of the NC induced by TF.
**(H)** Mass spectral envelopes corresponding to peptide 504-535 of β-gal $NC_{1\text{-}687}$, detected in HDX-MS data of isolated or TF bound (+TF) $RNC_{1\text{-}687}$ samples. The isotopic distributions were fit to single or bimodal gaussian curves (red and green) using HXExpressV3.

#### Figure S4

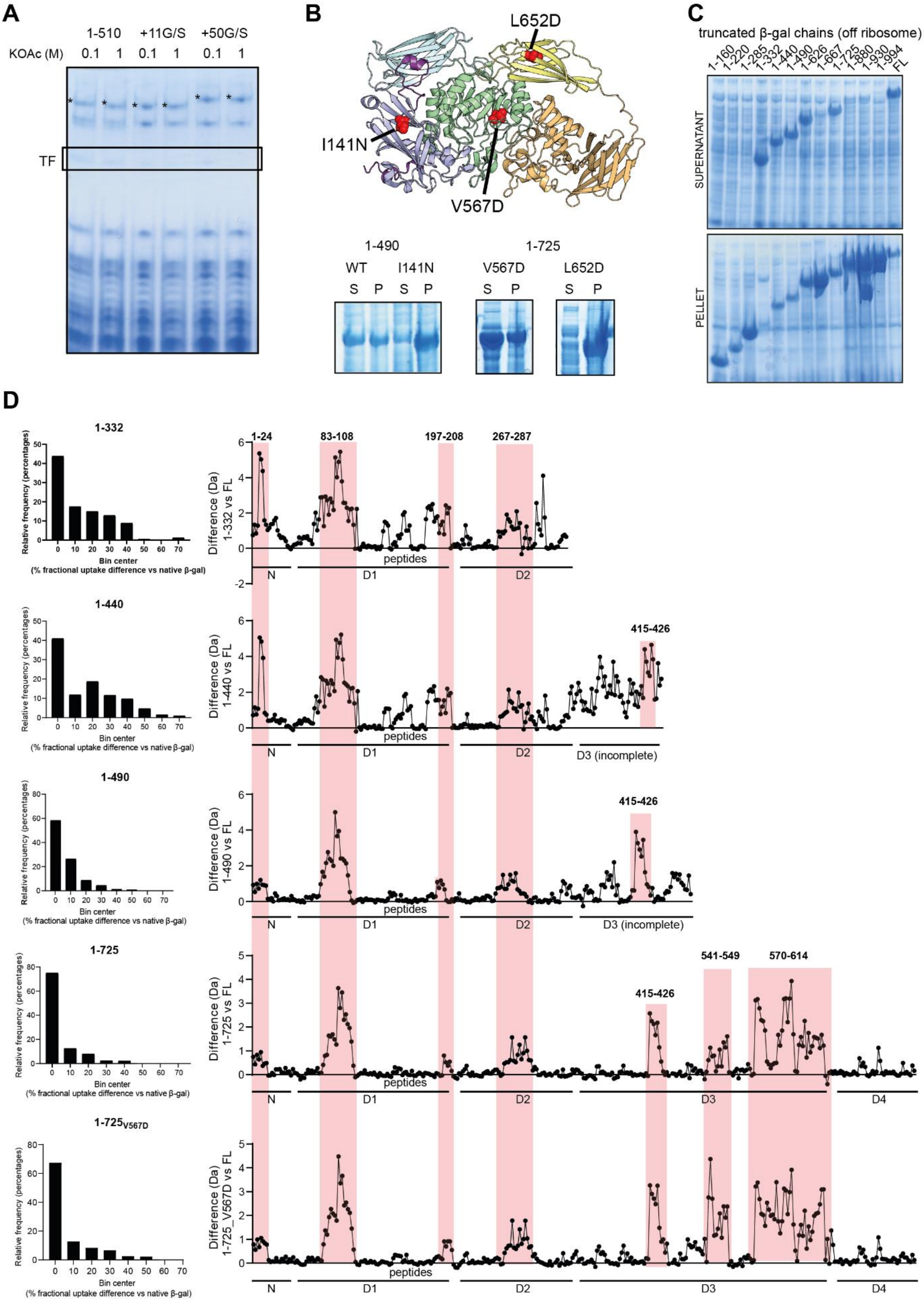

A
1-510
+11G/S
+50G/S
KOAc (M)
0.1 1 0.1 1 0.1 1
TF
B
L652D
I141N
V567D
1-490
1-725
WT
I141N
V567D
L652D
S P S P S P S P
C
truncated β-gal chains (off ribosome)
1-160 1-220 1-285 1-332 1-440 1-490 1-626 1-667 1-725 1-880 1-930 1-994 FL
SUPERNATANT
PELLET
D
1-332
1-440
1-490
1-725
1-725_V567D
Relative frequency (percentages)
Bin center
(% fractional uptake difference vs native β-gal)
Difference (Da) 1-332 vs FL
Difference (Da) 1-440 vs FL
Difference (Da) 1-490 vs FL
Difference (Da) 1-725 vs FL
Difference (Da) 1-725_V567D vs FL
1-24
83-108
197-208
267-287
415-426
541-549
570-614
peptides
N
D1
D2
D3 (incomplete)
D3
D4

**Figure S4. Effects of destabilising mutations and truncations on β-gal, related to Figure 4**

**(A)** Coomassie-stained SDS-PAGE of the resuspended ribosomal pellet from co-sedimentation assays of RNCs (1-510, 1-$510_{+11G/S}$ and 1-$510_{+50G/S}$) purified from WT *E. coli* and incubated with additional TF *in vitro*. The KOAc salt content in the sucrose cushion was 0.1 or 1 M as indicated. The expected position of TF is indicated, and bands corresponding to NCs are labelled (*).

**(B)** Top: Structure of β-gal (PDB: 6CVM) with residues I141, V567 and L652 shown as red spheres. Bottom: Coomassie-stained SDS-PAGE of the soluble (S) and pellet/insoluble (P) fraction of a cell lysate from *E. coli* cells overexpressing truncated β-gal constructs 1-$490_{WT}$, 1-$490_{I141N}$, 1-$725_{V567D}$ and 1-$725_{L652D}$.

**(C)** Coomassie-stained SDS-PAGE gels of the soluble (top, supernatant) and insoluble (bottom, pellet) fractions of a cell lysate from *E. coli* cells overexpressing truncated β-gal constructs. Full-length (FL) β-gal is included as a control in the last lane.

**(D)** Left: histograms showing the frequency distribution of residues based on the fractional deuterium uptake difference between truncated chains (1-332, 1-440, 1-490, 1-725 and 1-$725_{V567D}$) and full-length β-gal. Right: Plot of the difference in deuterium uptake (after 100 s deuteration) between truncated chains and full-length β-gal. Values are plotted for individual peptides covering β-gal in each chain, and regions that are destabilised in multiple chains are highlighted in red. Higher values indicate more deuteration of peptides in truncated chains relative to full-length β-gal. Uptake values are listed in Table S6.

#### Figure S5

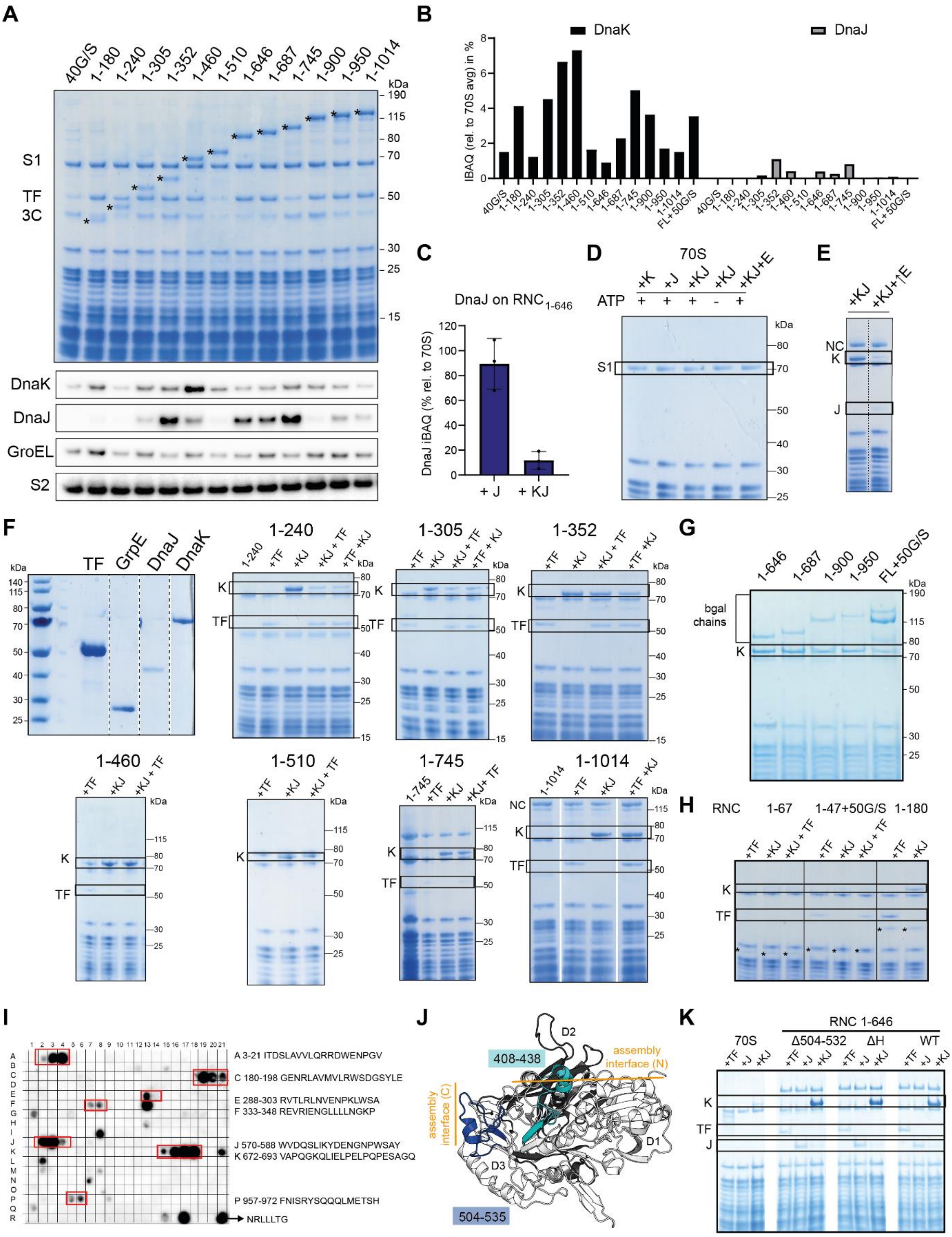
A
B
C
D
E
F
G
H
I
J
K
DnaK
DnaJ
GroEL
S2
S1
TF
3C
kDa
DnaJ on RNC 1-646
DnaJ iBAQ (% rel. to 70S)
+ J
+ KJ
iBAQ (rel. to 70S avg) in %
70S
ATP
NC
bgal chains
RNC
1-67
1-47+50G/S
1-180
A 3-21 ITDSLAVVLQRRDWENPGV
C 180-198 GENRLAVMVLRWSDGSYLE
E 288-303 RVTLRLNVENPKLWSA
F 333-348 REVRIENGLLLLNGKP
J 570-588 WVDQSLIKYDENGNPWSAY
K 672-693 VAPQGKQLIELPELPQPESAGQ
P 957-972 FNISRYSQQQLMETSH
NRLLLTG
D2
D1
D3
408-438
504-535
assembly interface (N)
assembly interface (C)
RNC 1-646
Δ504-532
ΔH
WT
K
J

**Figure S5. Determinants of DnaK/J binding to RNCs, related to Figure 5**

**(A)** Top: Coomassie-stained SDS-PAGE of β-gal RNCs purified via one 0.1 M KOAc sucrose cushion from WT BL21(DE3) *E. coli*. Bands corresponding to nascent chains (*) migrate slower than expected based on protein molecular weight, due to the covalently bound tRNA (~20 kDa). Some RNCs co-purify with Trigger factor (TF, ~50 kDa) at close to 1:1 stoichiometry. The band corresponding to 3C protease (~40 kDa) used in purification (Fig 1C) is indicated. All other major bands present in the purified RNC samples correspond to 70S ribosomal proteins including S1 (~70 kDa) Bottom: immunoblot of a replicate SDS-PAGE gel probed against DnaK, DnaJ, GroEL and ribosomal protein S2.

**(B)** Intensity-based absolute quantification (iBAQ) values for DnaK (black) and DnaJ (grey) normalised to the average iBAQ of all ribosomal proteins in each corresponding RNC sample, purified via one low-salt sucrose cushion.

**(C)** Mean iBAQ for DnaJ present in the resuspended ribosomal pellet from high-salt co-sedimentation assays of $RNC_{1\text{-}646}$, incubated with either DnaJ alone (+J) or with DnaJ and excess DnaK (+KJ), both in the presence of ATP. The iBAQ values were normalised to the average iBAQ value of all 70S ribosomal proteins. Error-bars correspond to the SD from 3 independent co-sedimentation assays.

**(D)** Coomassie-stained SDS-PAGE of the resuspended ribosomal pellet from high-salt co-sedimentation assays of 70S ribosomes incubated with either DnaK (+K), DnaJ (+J), or both DnaK and DnaJ (+KJ) in the presence and absence of ATP as indicated, as well as with additional GrpE (+KJ+E). The bands corresponding to ribosomal protein S1 are indicated.

**(E)** Coomassie-stained SDS-PAGE gel of the resuspended ribosomal pellet from high-salt co-sedimentation assays of $RNC_{1\text{-}646}$ incubated with either DnaK and DnaJ (+KJ) or with excess (25 μM) GrpE (+KJ+↑E), both in the presence of ATP. The bands corresponding to DnaK and DnaJ are indicated.

**(F)** Coomassie-stained SDS-PAGE of the resuspended ribosomal pellet from high-salt co-sedimentation assays of RNCs incubated, in the presence of ATP, with either TF (+TF), DnaK/DnaJ (+KJ), or all three factors added in the specified order (+KJ+TF or +TF+KJ). The bands corresponding to DnaK and TF are indicated. The first gel shows the migration of purified TF, GrpE, DnaJ and DnaK during SDS-PAGE, for reference.

**(G)** Coomassie-stained SDS-PAGE of the resuspended ribosomal pellet from high-salt co-sedimentation assays of RNCs incubated with DnaK and DnaJ in the presence of ATP. The bands corresponding to DnaK are indicated.

**(H)** Coomassie-stained SDS-PAGE of the resuspended ribosomal pellet from high-salt co-sedimentation assays of RNCs incubated with either TF (+TF) or DnaK with DnaJ (+KJ) or all three factors (+KJ+TF), all in the presence of ATP. The bands corresponding to DnaK and TF are indicated.

**(I)** Peptide array developed with ECL after incubation with DnaK and appropriate antibodies. Each position on the grid contains a 13-residue peptide derived from the wild-type β-gal sequence, starting from the initiator methionine (numbered as residue 0) at position A1. The sequence of peptides is shifted by 3 residues between neighbouring spots. The peptide in position R21 is the NRLLLTG peptide, serving as a positive control. Sites identified as potential DnaK binding sites are boxed in red and labelled with a letter (A, C, E, F, J, K, P) followed by the exact sequence.

#### Figure S6

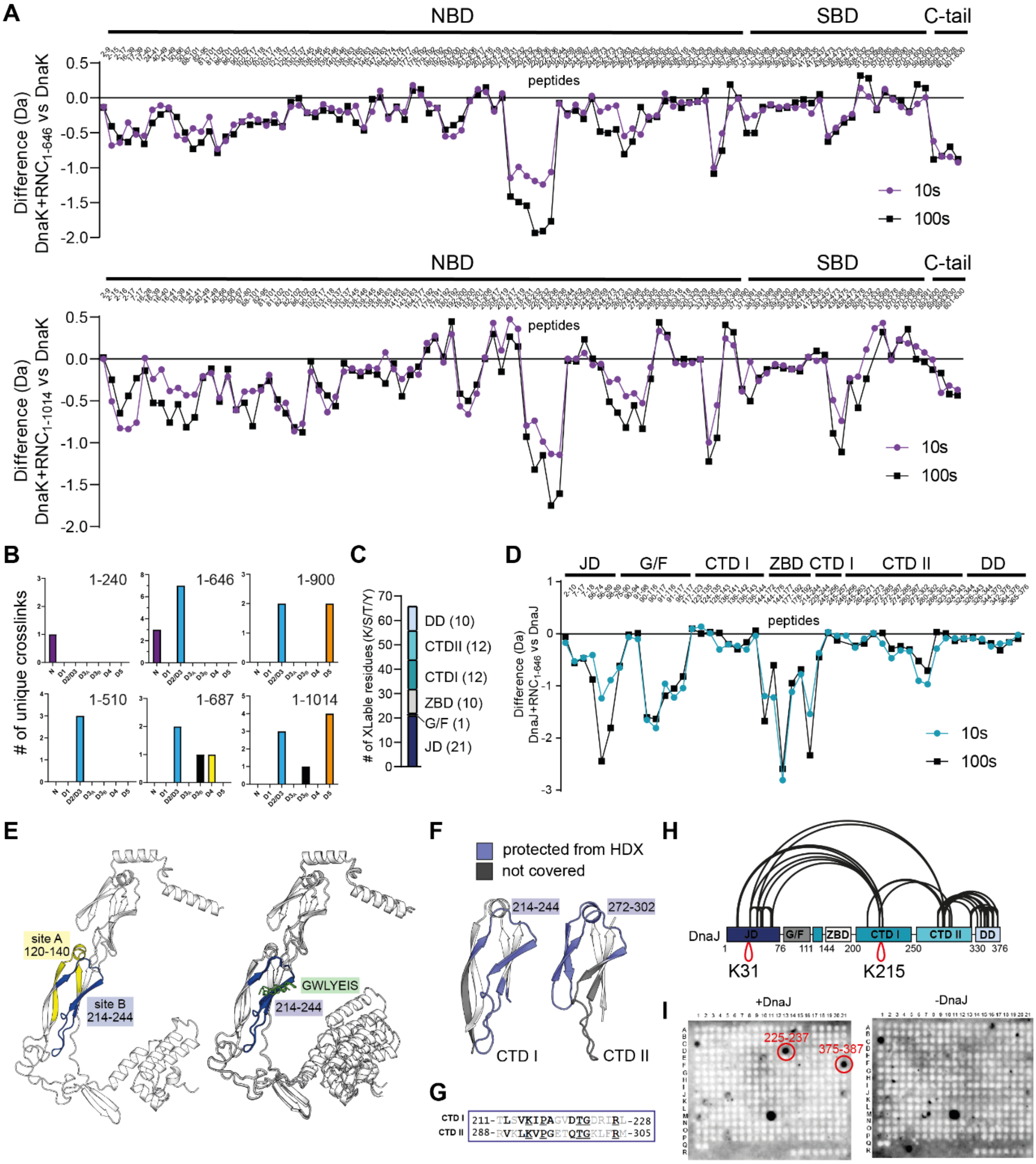


**Figure S6. DnaK/J interaction with nascent chains, related to Figure 5 and Figure 6**
**(A)** Difference in deuterium uptake after 10 s (purple) or 100 s (black) deuteration, between isolated DnaK and DnaK bound to $RNC_{1\text{-}646}$ (top) or $RNC_{1\text{-}1014}$ (bottom). Values are plotted for individual peptides covering DnaK detected in the HDX-MS dataset. Negative values indicate less deuteration of a peptide in RNC-bound DnaK relative to isolated DnaK. Uptake values are listed in Table S5.
**(B)** Number of unique crosslinks detected between DnaJ and each group of residues (as in Fig 3A) on the specified NC.
**(C)** Number of DSBU-reactive residues (K, S, T or Y) in each domain of DnaJ.

**(D)** Difference in deuterium uptake after 10 s (blue) or 100 s (black) deuteration, between isolated DnaJ and DnaJ bound to $RNC_{1-646}$. Values are plotted for individual peptides covering DnaJ detected in the HDX-MS dataset. Negative values indicate less deuteration of a peptide in RNC-bound DnaJ relative to isolated DnaJ.
**(E)** Left: DnaJ monomer structure (AF-P08622-F1), indicating site A (120-140, yellow) and site B (214-244, blue). Right: AlphaFold 2.0 multimer-predicted structure DnaJ monomer in complex with peptide GWLYEIS. The top 4 most confidently predicted structures are overlayed.
**(F)** Structure of CTD I (left) and CTD II (right) of DnaJ (AF-P08622-F1), with regions protected from deuterium uptake in RNC-bound DnaJ relative to isolated DnaJ coloured blue, and regions not covered by any detected peptides coloured grey.
**(G)** Sequence alignment of structurally homologous sites on CTD I and CTD II of DnaJ.
**(H)** Domain diagram of DnaJ showing intra-DnaJ crosslinks which were detected in DnaJ bound to $RNC_{1-646}$. Crosslinks between equivalent residues in two different DnaJ molecules - are shown in red.
**(I)** Peptide arrays developed with ECL after incubation with DnaJ and appropriate antibodies (left) or only with antibodies (right). Each position contains a 13-residue peptide derived from the wild-type β-gal sequence, starting from the initiator methionine (numbered as residue 0) at position A1. The sequence of peptides is shifted by 3 residues between neighbouring spots. Peptides identified as potential DnaJ binding sites are circled in red.

**Figure S7**

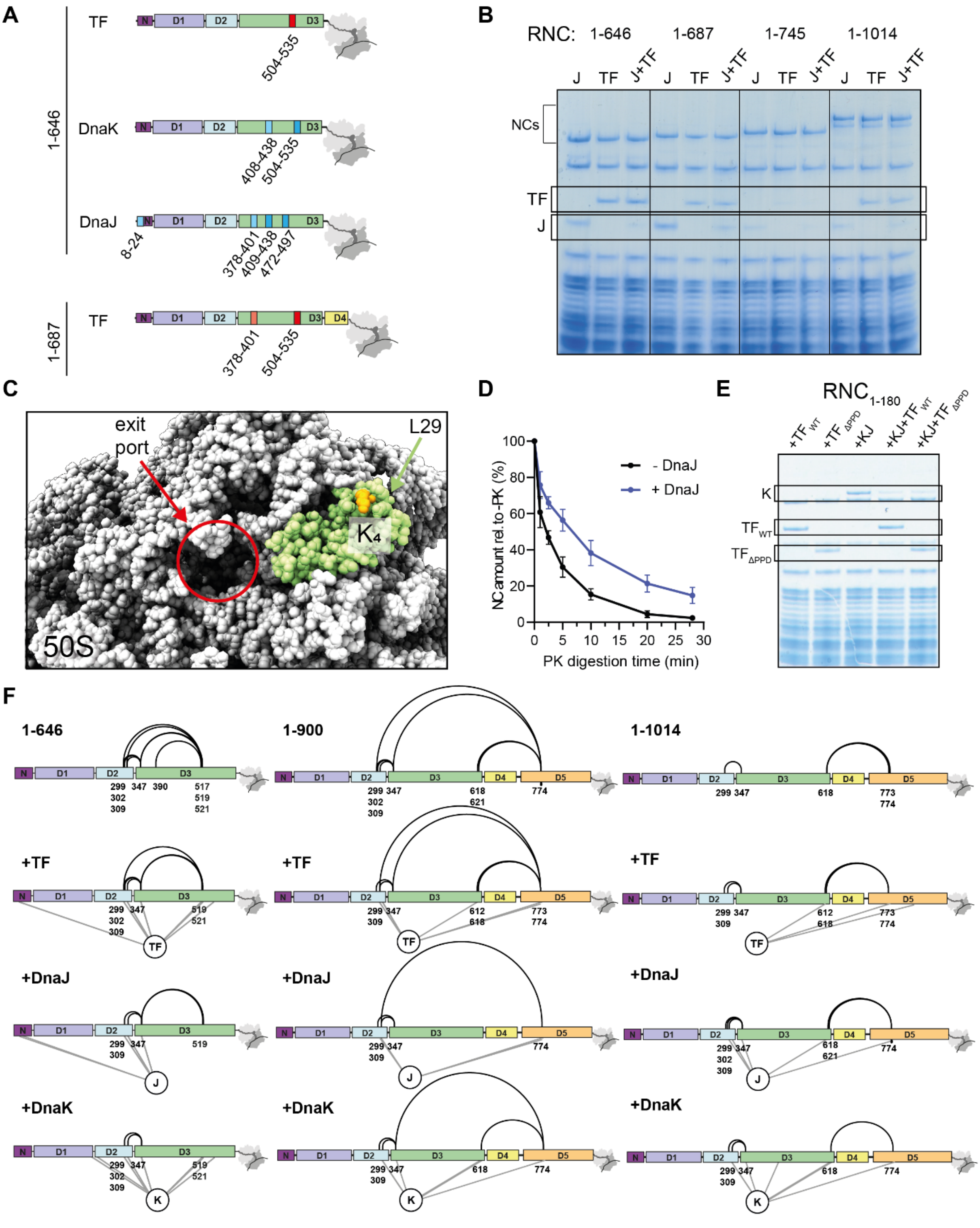


#### Figure S7. Cooperation and competition of DnaK, DnaJ and TF on RNCs, related to Figure7

**(A)** Domain schematic of $RNC_{1\text{-}646}$ (top) and $RNC_{1\text{-}687}$ (bottom), showing the position of sites affected by indicated chaperones based on HDX-MS experiments described in Fig 3, 5 and 6.

**(B)** Coomassie-stained SDS-PAGE of the resuspended ribosomal pellet from high-salt co-sedimentation assays of indicated RNCs incubated with either DnaJ (J), TF, or both (J+TF). The bands corresponding to DnaJ and TF are indicated.

**(C)** Close-up of the 70S ribosome exit port (red), with protein L29 coloured green and residue K4 in orange (PDB: 7D80). DnaJ, TF and β-gal nascent chains in RNC complexes formed crosslinks with a lysine on a tryptic peptide AKELR. Note that this peptide is present in both ribosomal proteins L29 ($_{3}$AKELR$_{7}$) and L17 ($_{41}$AKELR$_{45}$). However, only K4 in L29 is solvent-accessible.
**(D)** Amount of undigested $NC_{1\text{-}646}$ present at different time points after limited proteinase K digestion of $RNC_{1\text{-}646}$, in isolation (-DnaJ, black) or upon incubation with DnaJ (+DnaJ, blue). Error-bars correspond to the SD of 3 independent reactions.
**(E)** Coomassie-stained SDS-PAGE of the resuspended ribosomal pellet from high-salt co-sedimentation assays of $RNC_{1\text{-}180}$, incubated with either wild-type TF (+$TF_{WT}$), TF without the PPD (+$TF_{\Delta PPD}$), DnaK and DnaJ (+KJ) or combination of DnaK and DnaJ with either TF variant (+KJ+$TF_{WT}$ or +KJ+$TF_{\Delta PPD}$), all in the presence of ATP. The bands corresponding to DnaK and TF variants are indicated.
**(F)** Schematic showing the position of intra-NC (black) and chaperone-NC (grey) crosslinks detected in RNCs 1-646 (left), 1-900 (middle) and 1-1014 (right), with or without bound chaperones (TF, DnaJ or DnaK). Samples were prepared under four different conditions. From top to bottom: RNCs purified from ΔTF cells, RNCs purified from WT cells; RNCs purified from ΔTF cells and incubated with DnaJ; RNCs purified from ΔTF cells then incubated with DnaK, DnaJ and ATP. Crosslinked RNCs incubated with excess chaperones were subjected to an additional centrifugation through a high-salt sucrose cushion to remove unbound factors before trypsin digestion and MS analysis.

**Table S1.** Proteomic analysis of RNC composition in dataset A, related to Figure 1. List of detected proteins and corresponding intensities
**Table S2.** Proteomic analysis of RNC composition in dataset B, related to Figure 1. List of detected proteins and corresponding intensities
**Table S3.** Proteomic analysis of RNC composition in dataset C, related to Figure 1 and S5. List of detected proteins and corresponding intensities
**Table S4.** List of residues crosslinked in RNC samples purified from ΔTF or WT *E.coli*, related to Figure 2, 3 and 7
**Table S5.** Summary of differential HDX-MS of β-galactosidase RNCs with and without bound chaperones, related to Figure 2, 3, 5, 6 and 7
**Table S6.** Summary of differential HDX-MS of β-galactosidase truncations compared to native full-length β-galactosidase, related to Figure 4
**Table S7.** Proteomic analysis of resuspended pellets from co-sedimentation assays of RNCs with chaperones, related to Figure 5 and 7
**Table S8.** List of residues crosslinked in RNC samples purified from ΔTF *E. coli* and incubated with recombinant chaperones, related to Figure 5, 6 and 7
**Table S9.** LIMBO-predicted DnaK binding sites on β-galactosidase, related to Figure 5
**Table S10.** List of recombinant DNA used in this study, related to STAR Methods
**Table S11.** List of protein sequences of recombinant proteins used in this study, related to STAR Methods
